# Ketamine and Psilocybin Differentially Impact Sensory Learning During the Mismatch Negativity

**DOI:** 10.1101/2025.11.06.687023

**Authors:** Shona G. Allohverdi, Milad Soltanzadeh, André Schmidt, Colleen E. Charlton, Daniel J. Hauke, Povilas Karvelis, Franz X. Vollenweider, Andreea O. Diaconescu

**Author notes:** Corresponding author: Dr. Andreea Diaconescu. Equal contribution (Franz X. Vollenweider, Andreea O. Diaconescu).

## Abstract

Ketamine and psilocybin show potential as therapies for various mental illnesses, including major depressive disorder. However, further investigation into their neural mechanisms is required to understand their effects on the brain. By combining computational modelling with electroencephalography (EEG), we examine the effects of ketamine and psilocybin on hierarchical sensory pwPE learning in the context of the auditory mismatch negativity, an event-related potential consistently shown to be reduced under psychotomimetic interventions. We employed a Bayesian framework and re-analyzed a previously acquired EEG dataset (Schmidt et al., 2012) by modelling single-trial EEG data using the Hierarchical Gaussian Filter. Using a placebo-controlled within-subject crossover design, healthy subjects were administered either S-ketamine or psilocybin during an auditory roving paradigm of pure sinusoidal tones. Our findings elucidate distinct neural impacts of ketamine and psilocybin on sensory learning: ketamine led to a larger reduction in the effect of sensory precision compared to placebo from 207 to 316 ms peaking at 277 ms in the frontal central channels, while psilocybin showed no significant effect. Both drugs reduced the expression of belief precision between 160 to 184 ms, peaking at 172 ms. For higher-level volatility pwPEs, ketamine reduced the expression at 312 ms while psilocybin had a null effect. For perception of elementary imagery, ketamine had a greater effect than psilocybin on sensory and volatility precision, while psilocybin had a greater effect on volatility pwPEs. Our findings suggest hallucinogens have distinct effects on sensory learning that could inform tailored therapies for major depression.

## 1 Introduction

Psychedelics including psilocybin and dissociatives like ketamine are promising candidate treatments for various psychiatric disorders, particularly for major depressive disorder (MDD) [1]. N-methyl-D-aspartate receptor (NMDAR) antagonism by ketamine boosts glutamate release at *α*-amino-3-hydroxy5-methyl-4-isoxazolepropionic acid receptors (AMPAR), which leads to a series of events enhancing synaptic plasticity [2, 3]. Conversely, psilocybin primarily activates 5-hydroxytryptamine 2A (5-HT_2A_) receptors [4]. Both compounds have shown success in alleviating depression and anxiety symptoms [5, 6, 7, 8, 9, 10]. However, how these agents reduce symptoms remains poorly understood, warranting the necessity for more detailed mechanistic studies. Despite their distinct mechanisms of action, both ketamine and psilocybin induce psychotic-like experiences, including hallucinations, delusions and disorganized behaviour [11, 12]. Notably, ketamine can also cause negative symptoms of schizophrenia like anhedonia [11, 13, 14]. The underlying cognitive and neural mechanisms have been previously explored through electrophysiological studies using the auditory mismatch negativity (MMN) paradigm [15, 16, 17].

The auditory MMN, an electrophysiological signal elicited by violations of regularities in auditory input sequences, has been used as an empirical demonstration that the brain learns the statistical properties of its environment and anticipates upcoming sensory stimuli [18, 19, 20]. Predictive coding assumes that the auditory MMN reflects continuous model updating in a cortical hierarchy: each level generates predictions for the level below and receives prediction error (PE) signals when the sensory input diverges from those predictions [26, 27]. Predictions descend along backward connections, whereas PEs ascend along forward connections [28]. These pathways are thought to be neurochemically segregated: descending signals rely predominantly on NMDAR mediated transmission, whereas ascending signals rely more on AMPARs [29]. The influence of PEs is modulated by a learning rate. Operationally, this learning rate is the ratio of sensory precision (numerator: confidence in the bottom-up input) to belief precision (denominator: confidence in top-down priors [42]), where the precision corresponds to the inverse of the variance. Consequently, the MMN can be viewed as the difference between pwPEs elicited by surprising (‘deviant’) and predictable (‘standard’) events [27].

Physiologically, the MMN depends on NMDAR signalling. Ketamine administration, for instance, has been observed to reduce the MMN amplitude [21, 16, 22]. In contrast, psilocybin does not reduce MMN amplitudes [17, 15, 16], but has been observed to modulate other event-related potential (ERP) components associated with sensory learning such as the P300 [15, 16, 23]. For example, Bravermanova and colleagues [15] showed that psilocybin attenuates the P3, a component recently linked with high-level pwPEs [24]. Because the P3 reduction implies altered learning about environmental volatility, we predicted that psilocybin would modulate precision estimates that weight these PWPEs. Conversely, other findings suggests that the N1 reflects low-level sensory precision [25], which can increase the impact of sensory evidence relative to priors. In light of influential theories about the effects of psychedelics this led us to expect complementary psilocybin effects in early latency windows. Based on these findings, we hypothesized that psilocybin would impact both low- and high-level precision.

Predictive coding models therefore anticipate a cascade of hierarchically related pwPEs that unfold over time, and trial-wise fluctuations in ERP amplitude should mirror Bayesian belief updating [25, 24, 30]. Second, pwPEs should be sensitive to NMDAR manipulations, given that blocking NMDARs diminishes top-down (predictive) signaling, thereby reducing constraints on inference about sensory inputs and rendering predictable and less predictable stimuli more similar in their degree of surprise [14]. These ideas align with recent pharmacological evidence indicating that EEG signatures of hierarchical pwPEs in auditory oddball tasks are indeed sensitive to NMDAR antagonism [31] and cholinergic modulation [30].

Viewing the MMN as a pwPE difference waveform naturally motivates moving average ERPs to the temporal evolution of EEG amplitudes across trials. Trial-by-trial fluctuations in EEG amplitude carry information about how precision estimates—and the pwPEs they scale—are updated as the brain adapts to sensory irregularities [32, 24, 33, 34, 27, 35]. Investigating these dynamics therefore offers a window onto the mechanisms by which psychotomimetic and psychedelics such as ketamine and psilocybin reshape hierarchical learning.

### Study rationale and aims

We re-analyzed the placebo controlled cross over dataset of Schmidt et al. (2012) [16], in which healthy volunteers received S**-**ketamine and psilocybin during an auditory oddball task. The authors examined drug effects on the memory trace effect of the MMN [36]. The memory trace effect represents the increase in the MMN amplitude following an increase in the number of ‘standard’ tone repetitions. The study revealed a decline in the MMN slope at frontal channels under ketamine, but not psilocybin, which was interpreted as impaired auditory PE processing under ketamine. Later, Schmidt, Diaconescu and colleagues (2012) [16] modelled the effects of S-ketamine using dynamic causal modelling for event-related potentials [37, 38]. This study provided empirical evidence for the predictive coding framework of the MMN showing that it could be explained by changes in bottom-up and top-down effective connectivity, with ketamine reducing the bottom-up signalling from the primary auditory to the secondary auditory cortex; however, in this study, it was unclear whether the impact of ketamine on bottom-up effective connectivity reflected a reduction in PE signalling. Here, we directly assess the relationship between S-ketamine and belief precision and pwPEs by employing a hierarchical Bayesian model that allows us to model trial-by-trial belief updates under different pharmacological manipulations.

The relaxed beliefs under psychedelics (REBUS) model by Carhart-Harris and Friston [39], alongside Letheby and Gerrans’ self-binding model [40], suggest that psychedelics weaken high-level priors, potentially facilitating learning and belief revision. According to the REBUS model, psychedelics induce a state of heightened entropy in the brain, which may lead to a reduction in the precision of high-level predictions, thereby enhancing sensory learning. The ensuing increase in PE learning coupled with a decrease in the precision of higher-level predictions means that priors no longer constrain perceptual and cognitive processes, thereby facilitating new interpretations of sensory data leading to the generation of new ideas and perspectives [41].

Leveraging the placebo-controlled cross-over dataset of Schmidt et al. [16], we applied the Hierarchical Gaussian Filter (HGF)[42] to single trial EEG amplitudes to replicate the findings of [43] and quantify multiple, hierarchically linked precision estimates and pwPEs. This design enables formal testing of drug × hierarchical level interactions. We hypothesised that NMDAR antagonism with S-ketamine would blunt both low and high level pwPEs by reducing top-down precision, yielding a reduced correlation between model-derived pwPEs and EEG amplitudes, whereas 5 HT_2A_ agonism with psilocybin would shift the precision balance, thus diminishing high-level belief precision (manifested as attenuated late, P3 related PEs) while concurrently modulating low level sensory precision (indexed by early N1). We extend earlier dynamic causal modelling work [16, 31] by isolating the direct influence of precision signals, in addition to pwPE magnitude, on sensory learning, thereby clarifying how distinct neuromodulatory systems shape hierarchical inference in the human cortex.

## 2 Materials and Methods

For a detailed description of the protocol see [16]. Here, we summarize the pertinent details of the study protocol and the model-based EEG analysis.

### 2.1 Participants

39 participants (Ketamine group: N = 19; Psilocybin group: N = 20) were included in this study. Both groups did not differ significantly in weight, age or sex. For a detailed breakdown of inclusion criteria and questionnaire details see Supplementary Materials (see Inclusion Criteria Details section) and [16]. Four subjects in the psilocybin group were excluded from the analysis due to incomplete data (final sample: ketamine group: N=19 (male:13; mean age = 26 ± 5.36); psilocybin group: N=16 (male: 11; mean age = 23 ± 2.41, Table 1)). All participants gave informed written consent and the study was approved by the Ethics Committee of the University Hospital of Psychiatry, Zurich and the use of psychoactive drugs was approved by the Swiss Federal Health Office, Department of Pharmacology and Narcotics (Bern, Switzerland).

### 2.2 Experimental Procedure and Paradigm

Following confirmation of inclusion criteria (Supplementary 1.1) through medical assessment and questionnaires, participants were separated into two groups: S-ketamine or Psilocybin. In a double-blind, placebo-controlled within-subject crossover design, participants underwent two sessions (active drug (S-ketamine/psilocybin vs. placebos), with counterbalancing after two weeks. Participants remained under constant supervision until end-of-dose effects were observed, at which point, they were then released into the custody of a partner or relative. All participants confirmed they had not eaten breakfast or ingested caffeine prior to experimentation.

### S-Ketamine/Placebo condition

S-ketamine was administered using an indwelling catheter that was placed in the antecubital vein of the non-dominant arm. An initial bolus injection of 10 mg over 5 min was followed, after a 1 min break, by a continuous infusion with 0.006 mg/kg per min over 80 min. The initial dose was reduced by 10% every 10 min, titrating to maintain a constant S-ketamine plasma level. For the placebo condition, an identical procedure was followed using a physiological sodium chloride solution with 5% glucose. Recording of the auditory paradigm began 10 minutes after continuous infusion of ketamine.

### Psilocybin/Placebo condition

Psilocybin was obtained from the Swiss Federal Office for Public Health. Administration occurred via gelatin capsules at a dose of 115 *µ*g/kg per oz, informed by guidelines set in that of [44, 45] for psilocybin and an equal number of capsules containing placebo. Recording of the auditory paradigm began 90 minutes after administration of psilocybin. A psychometric examination using a revised version (see Supplementary 1.6) of the 5-Dimensional Altered States of Consciousness Questionnaire (5D-ASC) questionnaire was also administered 6-hours after experimentation to measure the subjective effects of ketamine and psilocybin on participants [46, 47]. EEG data was recorded during an auditory roving oddball paradigm [36, 31, 48] developed by [49]. Please see Supplementary material sections 1.2 and 1.3 for details on the paradigm and the visual distraction task, respectively.

#### 2.2.1 Data Processing

EEG was recorded using a 64 electrode BioSemi system at a sampling rate of 512 Hz. Pre-processing and data analysis was performed using SPM12 (v7771;(http://www.fil.ion.ucl.ac.uk/spm/) in MATLAB (R2021b). EEG recordings were referenced to the average, high-pass filtered using a Butterworth filter with a cutoff frequency 0.5 Hz, down-sampled to 256 Hz, and low-pass filtered using Butterworth filter with cutoff frequency 30 Hz. The data were epoched into 500 ms segments around tone onsets, using a pre-stimulus baseline of 100 ms.

Similar to [31], we employed an ocular movement artefact rejection procedure. All trials that overlapped with the eye blink events were rejected (see Table S2 and Supplementary section 1.4). An additional artefact rejection procedure was applied to remove any problematic trials or channels. Trials with amplitudes exceeding ±80 *µ*V in any channel relative to prestimulus baseline were removed.

### 2.3 Computational Model

To model participants’ learning, we employed a multivariate version of the HGF [31, 43]. The HGF is a generic Bayesian model and was applied to model learning in various contexts, including associative learning [51], reward learning [52] and the MMN [31, 30, 33, 24].

For additional details about the model, see Weber et al., (2020) [31]. Scalars are denoted by lower case italics, vectors by lower case bold font, and matrices by upper case bold font. This model assumes that an agent infers two continuous quantities of hidden states during the auditory roving paradigm: The multivariate transition contingencies from one of the seven tone frequencies to the next denoted by the matrix **X**_2_, and the common volatility amongst these transitions, or how quickly contingencies change over time, is denoted by *x*_3_. On each trial *k*, the elements of **X**_2_ determine the probability of encountering each of the possible transitions **X**_1_, which in turn give rise to a particular tone **u** being heard or trial outcome **u**. The evolution of the elements of the transition matrix **X**_2_ are determined by Gaussian random walks, where the step-size is a function of the current value of the volatility state *x*_3_, based on the assumption that agents learn faster under a changing compared to a stable environment. An illustration (inspired by [31]) of the hierarchical states and their dependencies are shown in Figure 1.

**Figure 1:**
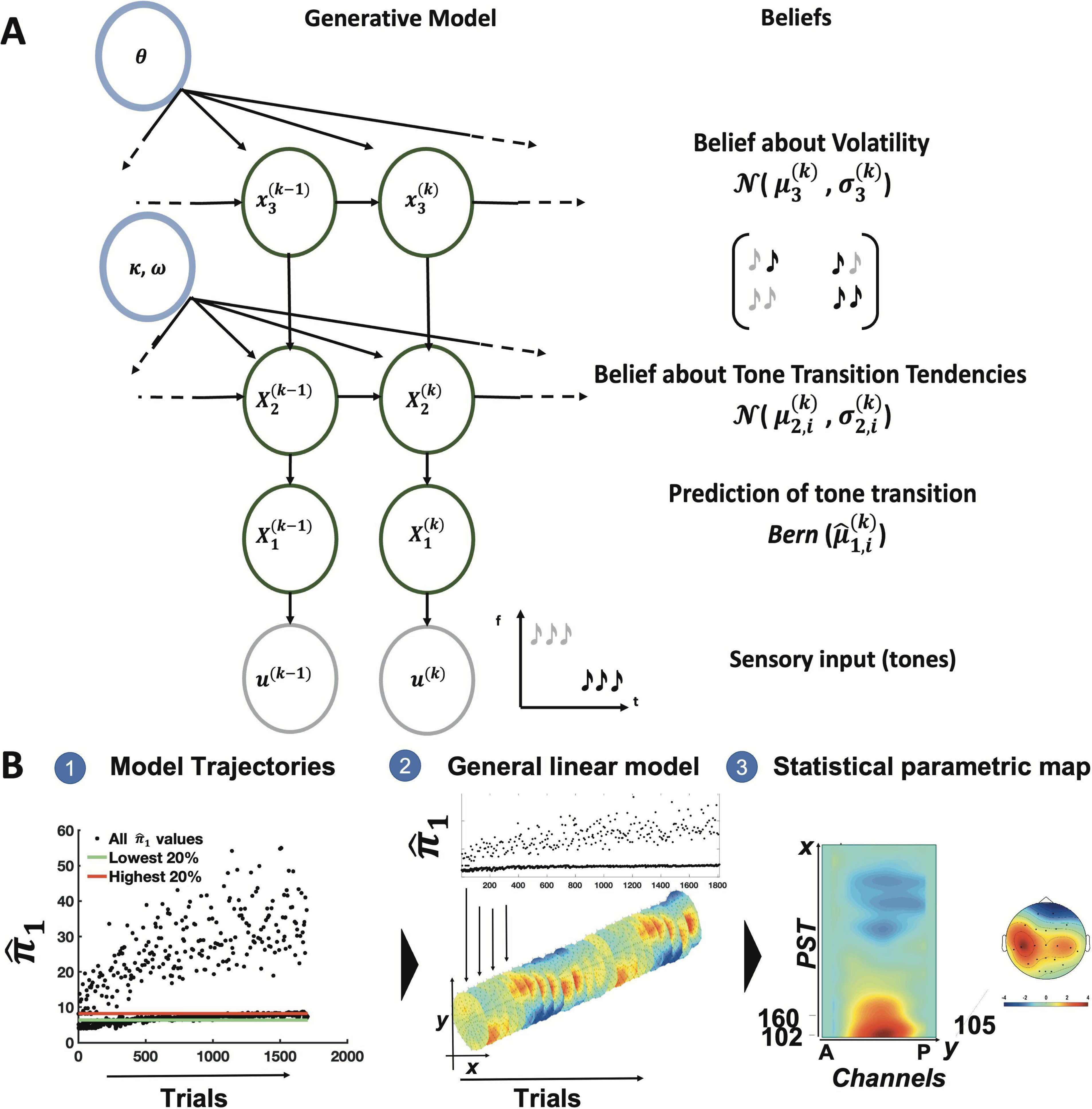
The Computational Model: A multivariate version of the binary three-level Hierarchical Gaussian Filter (HGF) **(A)**, previously employed in Within this schema, the agent is tasked with the inference of two continuous dimensions: the transition propensities between distinct tonal frequencies, encapsulated within the transition matrix **X**_2_, and a unified metric of the volatility of these transition propensities, symbolized as *x*_3_. To employ this model, the agent’s beliefs are updated via simple one-step update rules. Beliefs are mathematically characterized by their mean *µ* and variance *σ*. **Computational Single-Trial Analysis (B)**: (**1**) Simulated optimal Bayesian belief trajectories for each parameter of interest (π^_1_^(*k*)^shown in example) were used as predictors in a General Linear Model (GLM) to explain amplitude changes across trials in the single trial EEG data (**2**), to establish general linear models (GLMs).

The three parameters governing this model, *κ*, *ω*, and *ϑ*, determine the strength of the coupling between the second and third level, the constant learning rate at the second level that is independent of the third level, and the variance of the volatility over time (meta-volatility), respectively.

Since the MMN paradigm is a passive task that does not require participants to make responses, we optimized the parameters of the perceptual model assuming an ideal Bayesian observer that minimizes the cumulative Shannon surprise for a given input sequence using the ‘tapas_bayes_optimal binary_transition’ function from the TAPAS toolbox (v3.0; open-source code available as part of the TAPAS software collection:[53], https://www.translationalneuromodeling.org/tapas). Perceptual and prior parameter estimates are summarized in Supplementary Table 2. For the current analysis, we used the default priors in the HGF toolbox which include *µ*_2_ *µ*_3_, *σ*_2_,*σ*_3_, *ω*, *θ*, and *κ* as specified in the tapas_hgf_transition_config (see Supplementary Table S3).

#### 2.3.1 Inverting the Model: The Update Equations

The variational inversion of the above generative model under a mean-field approximation yields simple one-step update equations that, collectively, represent a perceptual model in which beliefs are updated via pwPEs or ε_i_^(*k*)^ [43] at hierarchical level *i* and on trial *k* (Eq.1). “Belief” refers to a posterior probability distribution as described by its sufficient statistics (assuming Gaussian distributions), i.e., mean *µ* and variance *σ* (or its inverse representative, the precision *π*). Predictions and precisions of predictions, i.e., prior precisions about the hidden states (before hearing the tone), are denoted with a hat symbol (e.g., π^).

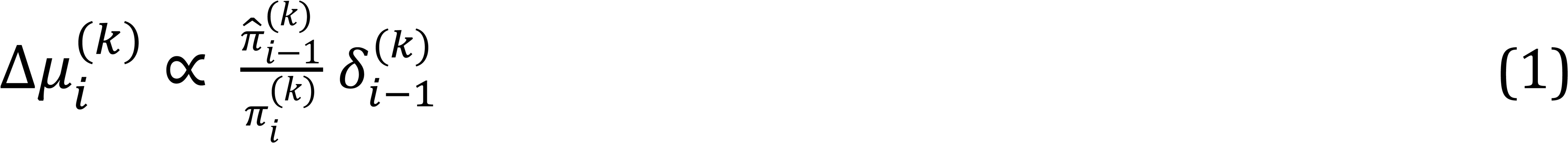

Wherein:

- μ*_i_*^(*k*)^denotes the posterior mean of the belief on each trial *k* of level *i* in the hierarchy.
- π^*_i-1_*^(*k*)^ is the precision of the prediction from the level below
- π*_i_*^(*k*)^ is the precision of the belief at the current level of the hierarchy
- δ*_i-1_*^(*k*)^ is the PE about the state of the level below.

#### 2.3.2 Computational Quantities Related to the MMN

The MMN is assumed to reflect hierarchically-coupled pwPEs in the auditory sensory stream [24, 31]. pwPEs are comprised of δ^(*k*)^, which denotes the PE about the level below, weighted by a precision ratio, (π^^(*k*)^) the predicted sensory precision at the level below, and (π^(*k*)^) representing the precision of the prediction at the current level (the belief precision).

The intuition behind this is that we place a greater emphasis on PEs when lower-level (sensory) inputs are precise compared to the precision of higher-level predictions. While low-level sensory pwPEs were related to the MMN, high-level volatility pwPE were strongly correlated with the P300 component [24].

#### 2.3.3 Statistical Analysis

The experimental pipeline is shown in Figure 2. We simulated optimal belief trajectories for ε_2_^(*k*)^, ε_3_^(*k*)^, π^_1_^(*k*)^, π^_2_^(*k*)^ and π^_3_^(*k*)^. These computational variables informed a GLM of the trial-by-trial amplitude fluctuations for each subject. Statistical analyses were constrained to 100 - 400 ms peri-stimulus time (PST) in line with previous analyses [31, 30, 33, 24]. Our GLMs including subject-specific trial-wise pwPEs and precisions derived from the HGF tested the null hypothesis that the ensuing correlation with EEG amplitude is zero at each sensor and PST. This produced a statistical parametric map of the F-statistic reporting the significance of the F-statistic allowing us to assess *where* in sensor-space and *when* in time, correlations with pwPEs (or precisions) were expressed (see Supplementary section 1.5 for details).

**Figure 2:**
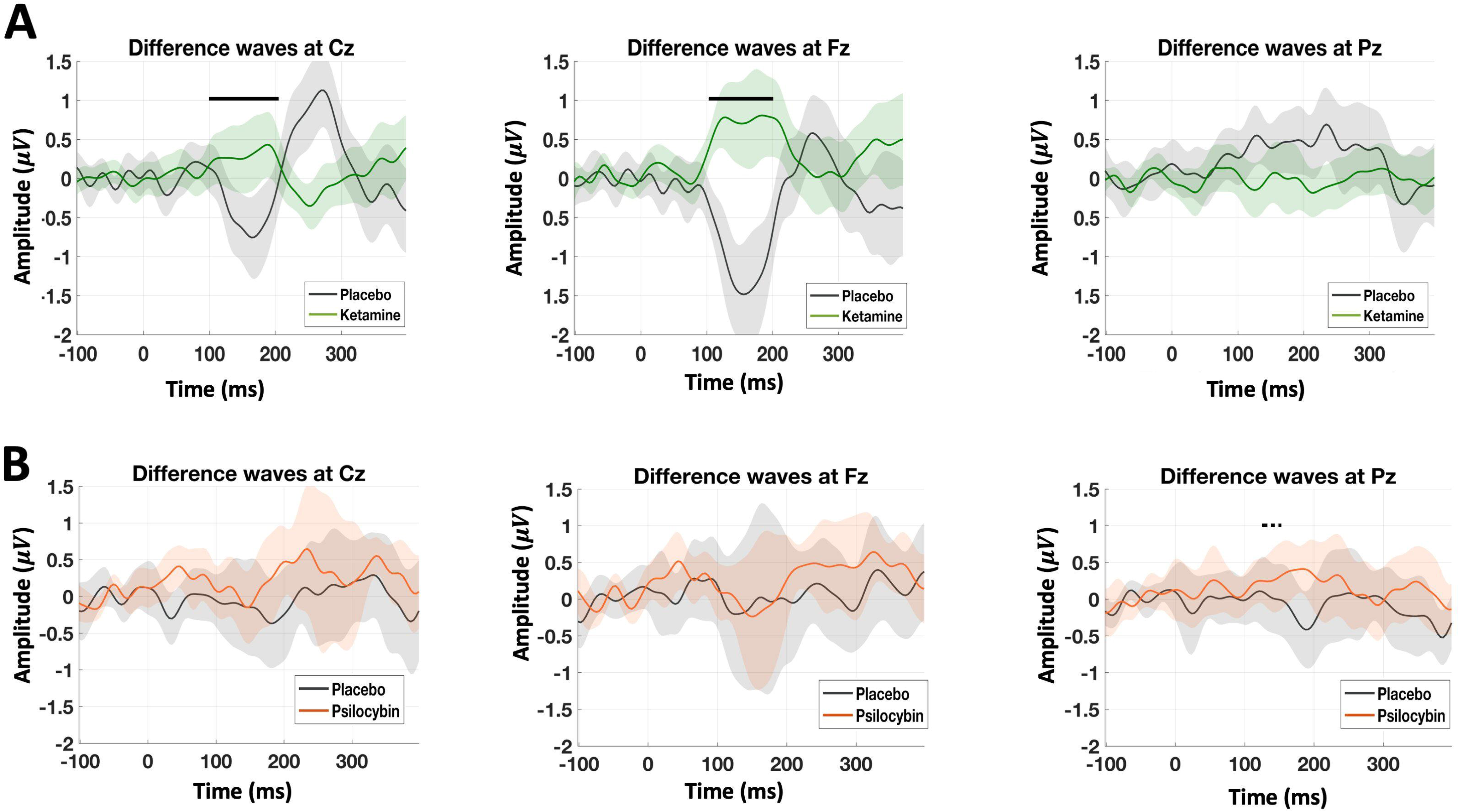
Grand-average MMN waveforms: (A) Ketamine (green) versus placebo (grey). (B) Psilocybin (orange) versus placebo (grey). Traces depict deviant–standard difference waves (mean ± SEM) at Cz, Fz, and Pz. Horizontal black lines mark intervals that reach significance after whole volume FWE correction (p < 0.05); these are present only in the ketamine comparison **(placebo > ketamine: *T*_(1,18)_ = −5.16, p *<* 0.00001, sensor: Cz, Fz)**. For psilocybin, we found trend level effects in posterior channels indicated by the dashed black lines **(peak: 145 ms, *T* = 5.35*, p_peak_* = 0.085)** with a cluster-defining threshold of p *<* 0.001

At the group level, we applied a 2 x 2 factorial design with within-subject factor ‘condition’ (active drug vs placebo condition) and between-subject factor ‘drug’ (Group 1: Ketamine vs Placebo vs Group 2: Psilocybin vs Placebo). We ran separate GLMs using this factorial design for each computational regressor and performed post-hoc t-tests to unpack the drug-by-condition interaction effects (Supplementary Section 1.5). We report the group-level results for each parameter separately, with the main effect of drug, study/type of pharmacological agent, and the interaction between drug and condition. Additionally, to test whether drug-induced alterations in precision and pwPE expression covary with subjective experiences, we implemented an ANCOVA including each participant’s 5D-ASC scores from the active sessions as covariates. We focused on the second-order 5D-ASC dimensions previously shown to differentiate ketamine from psilocybin, including disembodiment (DE), impaired control and cognition (ICC), and elementary imagery (EI). All statistical maps were corrected across sensors and timepoints for multiple comparisons using whole-volume family-wise error (FWE) control: peak-level and cluster-level FWE p *<* 0.05 (cluster defining threshold p*<* 0.001) [54, 55], using Gaussian Random Field theory [56].

#### 2.3.4 Code Availability

The analysis code for this study is publicly available at https://github.com/G-Verdi/MnketAnalysis.

## 3 Results

To enhance readability, results are presented in three nested subsections that proceed from descriptive, model-free contrasts to increasingly mechanistic analyses: (i) model-free comparisons of conventional mismatch negativity (MMN) difference waves across drug and condition; (ii) binned ERP analyses contrasting “high” versus “low” pseudo-conditions derived from the upper and lower quantiles of trial-wise computational variables (belief precisions and their pwPEs; Supplementary material); and (iii) parametric single-trial analyses in which trial-wise amplitude fluctuations are modeled by Hierarchical Gaussian Filter (HGF)–derived computations (three precisions of prediction and two pwPEs) to probe learning on a trial-by-trial basis.

### 1. Model-free analysis: Mismatch Negativity

Replicating prior work showing that NMDAR blockade diminishes auditory prediction-error signals (Schmidt et al., 2012), ketamine significantly attenuated MMN amplitude within the canonical 100–200 ms window over frontal midline sites (placebo > ketamine: *T*_(1,18)_ = −5.16, *p <* 0.00001, sensor: Fz and Cz). Psilocybin did not produce a statistically significant change in MMN amplitude across Fz and Cz channels (all p > 0.10). There was, however, a relative increase across parietal electrodes (peak = 145 ms, *T*_(1,15)_ = 5.58, *p_peak_*= 0.085, sensor: Pz), but this effect did not survive correction and is interpreted cautiously owing to the limited sample size (n= 16).

### 2. Binned ERP Analyses

To isolate condition-specific cortical responses, we adopted a quantile-binning approach. For each participant, trials were ranked by computational quantities derived from the three-level HGF. We then averaged ERPs for the lowest 20% and highest 20% of trials and entered the resulting bin-specific waveforms into a 2 × 2 mixed model (Drug × Computational Quantity Bin). Overall, this analysis showed that ketamine markedly attenuated prediction precision and error signaling, whereas psilocybin’s effects were confined to the sensory-level precision.

### Precision of Predictions

#### Sensory Precision (π^^(*k*)^)

Under ketamine, the effect of sensory precision was substantially reduced relative to placebo (peak = 281 ms, *T*_(1,18)_ = 5.00, *p_clus_*= 0.0001). By contrast, psilocybin induced an increase of sensory precision-related responses over central channels between 150 and 300ms (peak = 156 ms, *T*_(1,15)_ = 6.70, *p_peak_*= 0.028). Thus, ketamine broadly diminished, whereas psilocybin selectively augmented, trial-binned sensory-precision signals at 281 and 150-300ms ms, respectively (Figure S1 and Supplementary Table 5).

#### Belief Precision & Volatility Precision (π^^(*k*)^_2_, π^^(*k*)^_3_)

Neither drug produced statistically significant modulations of trial-binned belief and volatility precision (Supplementary Table 5).

### Precision-weighted PEs

#### Outcome Precision-Weighted PE (ε^(*k*)^_3_)

Overall, neither drug significantly modulated trial-binned outcome-level pwPEs after correction for multiple comparisons. Ketamine produced only a trend-level attenuation of outcome pwPE signals across a broad fronto-central cluster (peak = 223 ms, *T*_(1,18)_ = 4.98, *p_clus_*= 0.07). Psilocybin elicited no suprathreshold changes in either direction. (Supplementary Table 5).

#### Volatility Precision-Weighted PE (ε^(*k*)^_3_)

Ketamine elicited a fronto-central reduction in high-level pwPE, reaching cluster significance (peak = 168 ms, *T*_(1,18)_ = 5.05, *p_clus_*= 0.04). In contrast, psilocybin did not produce any suprathreshold modulation in either direction (Figure S2 and Supplementary Table 5).

### 3. Model-based Analyses

Across drugs and placebo conditions, all five computational quantities, three precisions of predictions and two pwPEs, showed robust correlations with trial-wise scalp EEG in epochs overlapping with classical mismatch-negativity effects, each occupying partially overlapping spatio-temporal effects. Precision signals (π^) dominated frontocentral sites, emerging early and reappearing later, whereas pwPEs (ε) involved broader centro-parietal regions and extend further in time. The temporal cascade, from early sensory certainty (π^_1_^(*k*)^, π^_2_^(*k*)^) to successive low- and high-level error signals (ε_2_^(*k*)^, ε_3_^(*k*)^), aligns with hierarchical predictive-processing accounts, in which predictions related to tone and sequence precision precede the processing of belief updates in the form of pwPE. All main effects survive whole-volume family-wise error correction (cluster-level *p* < 0.05, cluster-defining *p* < 0.001), with peak-level-corrected maxima highlighted in the figure (Figure 3, Figure S3, and Supplementary Tables 6 and 7).

**Figure 3:**
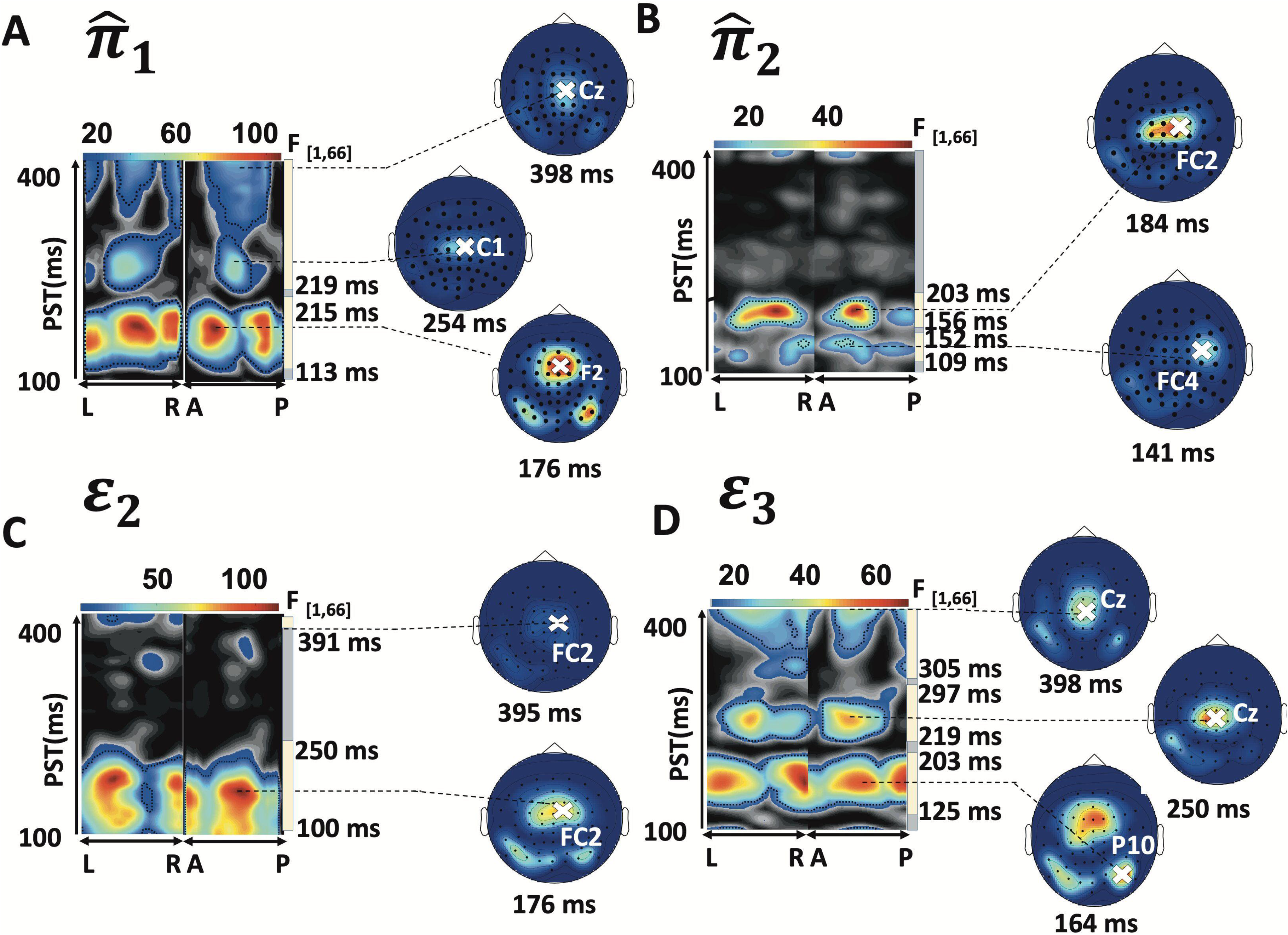
Main effects of HGF model parameters across conditions. Belief precision (π^_2_^(*k*)^) sensory precision (π^_2_^(*k*)^), low-level pwPE (ε_3_^(*k*)^), and high-level pwPE (ε_3_^(*k*)^)The topographical maps show maximum intensity projections for significant clusters of activation across space ((L) left to (R) right and (A) anterior to (P) posterior) and peristimulus time (PST), while the scalp maps show the activation corresponding to the peak of each cluster. Time windows of significant correlations are highlighted by the vertical yellow bars. Jet colour mapping displays cluster-level effects (p*<*0.05, whole-volume family-wise error (*FWE*) corrected at the cluster level with a cluster-defining threshold of p*<*0.001) while significant peak-level effects (p*<*0.05, whole-volume FWE-corrected at the peak level) are denoted with black dotted contours.

### Precision of Predictions

#### Sensory Precision (π^^(*k*)^_2_)

Using a full factorial design, we tested for drug-by-condition interaction effects, and found no significant interactions for π^^(*k*)^(peak = 309 ms, *F*_(1,66)_ = 13.43, *p_clus_*= 0.476, sensor: C3). We found a significant main effect across both condition and drug for π^^(*k*)^in the frontal and central channels (cluster 1: 113 - 215 ms, peak: 176ms, *F*_(1,66)_ = 128.54, *p_peak_* = 0.001; cluster 2: 219 - 400ms, peak: 254ms, *F*_(1,66)_ = 61.05, *p_peak_* = 4.88 E-07; cluster 3: 316 - 400 ms, peak: 398ms, *F*_(1,66)_ = 43.01, *p_peak_* = 5.13*E*−05) (Figure 3 A, Supplementary: Table S6).

Through exploratory post-hoc tests, we observed a reduction in the correlation between π^^(*k*)^trajectories derived from the ideal Bayesian observer and EEG amplitudes in ketamine compared to placebo (placebo *>* ketamine) (peak: 277 ms, *T*_(1,66)_ = 3.84, *p_clus_* = 0.003, sensor: C4; Figure 4A). Similar to the binned ERP analysis (Supplement), the effect of π^^(*k*)^ was significantly reduced under ketamine, compared to placebo.

**Figure 4:**
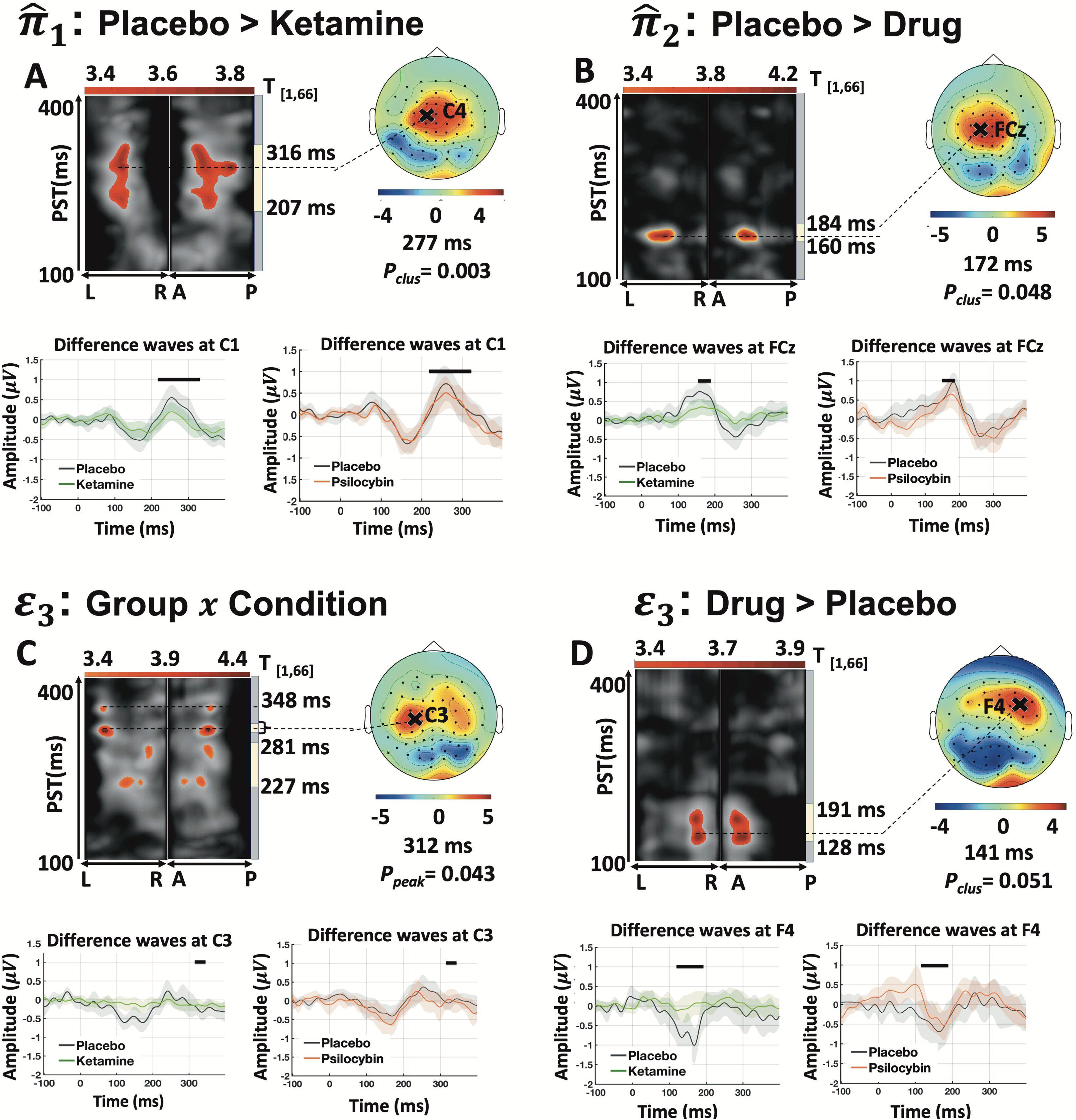
Main effects of HGF model parameters split by drug condition. (A) Sensory precision (π^_1_^(*k*)^) (B) Belief precision (π^_2_^(*k*)^) for (C) and (D) High-level precision-weighted prediction errors (pwPEs) (ε_3_^(*k*)^) *Top*: Maximum intensity projections of significant clusters across space ((L) left to (R) right and (A) anterior to (P) posterior) and peri-stimulus time (PST) are shown as topographical maps. Cluster extent is highlighted by the vertical yellow bar. Jet colour mapping displays cluster-level effects (p*<*0.05, whole-volume family wise error (*FWE*) corrected at the cluster level with a cluster-defining threshold of p*<*0.001) while significant peak effects (p*<*0.05, whole volume FWE-corrected at the peak level) are denoted with black dotted contours. Scalp maps to the right show cluster effects in jet colour map across a 2-D sensor layout. *Bottom*: ERP difference waveforms averaged for the 20 % highest and 20% lowest prediction error/precision values for ketamine (left) and psilocybin (right) conditions at electrodes in the vicinity of the global maximum. The time windows of significant effects from the parametric analysis are indicated by black bars. Please note that these ERP plots are merely meant to give an intuition of the effects; the actual statistical tests were performed jointly on all channels using the computational quantities as parametric regressors.

#### Belief Precision (π^^(*k*)^_2_)

No drug × condition interactions were detected (peak 391 ms, *F*_(1,66)_ = 12.23 *p_clus_* = 0.589, sensor FT7). Main effects across both condition and drug appeared at fronto-central channels (cluster 1: 109–152 ms, peak 141 ms, *F*_(1,66)_ = 22.13, *p_peak_* = 0.030; cluster 2: 156–203 ms, peak 184 ms, *F*_(1,66)_ = 54.44, *p_peak_* = 2.69 × 10⁻⁶). Furthermore, a main effect of drug was present between 160–180 ms (peak 172 ms, *T*_(1,66)_ = 4.30, *p_clus_* = 0.048, sensor FCz; Fig. 4B), driven by reduced placebo > ketamine correlations (peak 176 ms, *T_(_*_1,66)_ = 4.38, *p_clus_* = 0.043, sensor C1).

#### Volatility Precision (π^^(*k*)^_2_)

No significant Drug × Study interactions were detected. Main effects across both condition and drug were observed at fronto-central and posterior electrodes (cluster 1: 102–144 ms, peak 102 ms, *F_(_*_1,66)_ = 59.55, *p_peak_* = 8.90 × 10⁻⁷; cluster 2: 102–152 ms, peak 113 ms, *F*_(1,66)_ = 58.87, *p_peak_* = 1.05 × 10⁻⁶).

### Precision-weighted PEs

#### Low-level Precision-Weighted PEs (ε^(*k*)^_3_)

Main effects are shown in Figure 3C and Table S7. Neither Drug × Condition interaction (peak 375 ms, *F*_(1,66)_ = 16.41, *p_clus_* = 0.322, sensor F6) nor main effects of drug (ketamine: peak 375 ms, *F*_(1,66)_ = 12.83, *p_clus_* = 0.539, sensor F6; psilocybin: no surviving clusters) were observed, consistent with [40].

#### High-level Precision-Weighted PEs (ε^(*k*)^_3_)

A drug × condition interaction emerged between 312–320 ms over central channels (peak 312 ms, *T*_(1,66)_ = 4.33, *p_peak_*= 0.043, sensor C3; Fig. 4C). Post-hoc tests revealed reduced correlations under ketamine versus placebo (peak 223 ms, *F*_(1,66)_ = 23.83, *p_clus_* = 0.009, sensor FCz; *T*_(1,66)_ = 4.88, *p_peak_* = 0.026), with no analogous psilocybin effect. Main effects are shown in Figure 3D and Table S7. We also observed a marginal drug main effect over frontal channels (128–191 ms; peak 141 ms, *T*_(1,66)_ = 3.92, *p_clus_* = 0.051 FWE-corrected, sensor F4), driven by ketamine-related attenuation (Figure 4D). Within the ketamine group alone, the correlation with ε^(*k*)^ was significantly reduced by the drug (peak 223 ms, *T*_(1,18)_ = 5.92, *p_peak_* = 0.026, sensor FCz), consistent with [40]. Difference-image analyses (placebo–drug) further confirmed stronger ketamine-than-psilocybin reductions ((placebo–ketamine) vs (placebo–psilocybin); peak 230 ms, *F*_(1,27)_ = 22.87, *p_clus_* = 0.044; post-hoc *T*_(1,27)_ = 4.78, *p_clus_* = 0.021, *p_peak_* = 0.043; sensor FCz). These results confirm that relative to placebo, we see a significant reduction in correlations between ε^(*k*)^derived from the ideal Bayesian observer and EEG amplitudes in the ketamine group compared to psilocybin.

### 4. Correlations with Altered States Questionnaire Self-reports Precision of Predictions

After accounting for covariates, an omnibus F-test revealed pronounced drug differences for π^^(*k*)^ over posterior channels (peak = 355 ms, *F*_(1,27)_ = 31.65, *p_clus_* = 0.012, *p_peak_* = 0.020, sensor: P7). Post-hoc two-sample t-tests revealed a significant reduction in the correlation between Bayes optimal π^^(*k*)^ trajectories and EEG amplitudes for ketamine relative to psilocybin (peak = 355 ms, *T*_(1,27)_ = 5.63, *p_clus_* = 0.011, *p_peak_* = 0.010, sensor: P7).

### Associations with Subjective Experiences

#### Elementary Imagery

No significant drug differences were detected for π^^(*k*)^ (peak 363 ms, *F*_(1,27)_ = 18.21, p*_clus_* = 0.445, sensor FP2) or π^_1_^(*k*)^ (no surviving clusters). By contrast, π^_3_^(*k*)^ showed a significant drug difference (peak 395 ms, F_(1,27)_ = 35.16, p*_peak_* = 0.013, sensor T7), driven by a larger positive correlation under ketamine (peak 395 ms, F_(1,27)_ = 81.48, *p_peak_* < 0.001, sensor T7; psilocybin: no surviving clusters). A positive correlation between the effect of π^^(*k*)^ and elementary imagery at left frontal sites was observed across participants (peak 391 ms, T_(1,27)_ = 4.37, *p_peak_* = 0.037; Figure 5B), but this did not survive removal of one outlier (see Supplement). Additionally, across both drugs, the effect of π^^(*k*)^ correlated positively with imagery at right-frontal FC6 (∼121 ms; peak 121 ms, T_(1,27)_ = 5.45, *p_clus_* = 0.045, *p_peak_* = 0.014), an effect largely driven by ketamine (peak 121 ms, T(1,27) = 5.38, *p_peak_* = 0.017). In conclusion, ketamine enhances associations between both sensory (∼121 ms) and volatility (∼395 ms) precision signals and elementary imagery, whereas psilocybin does not.

**Figure 5:**
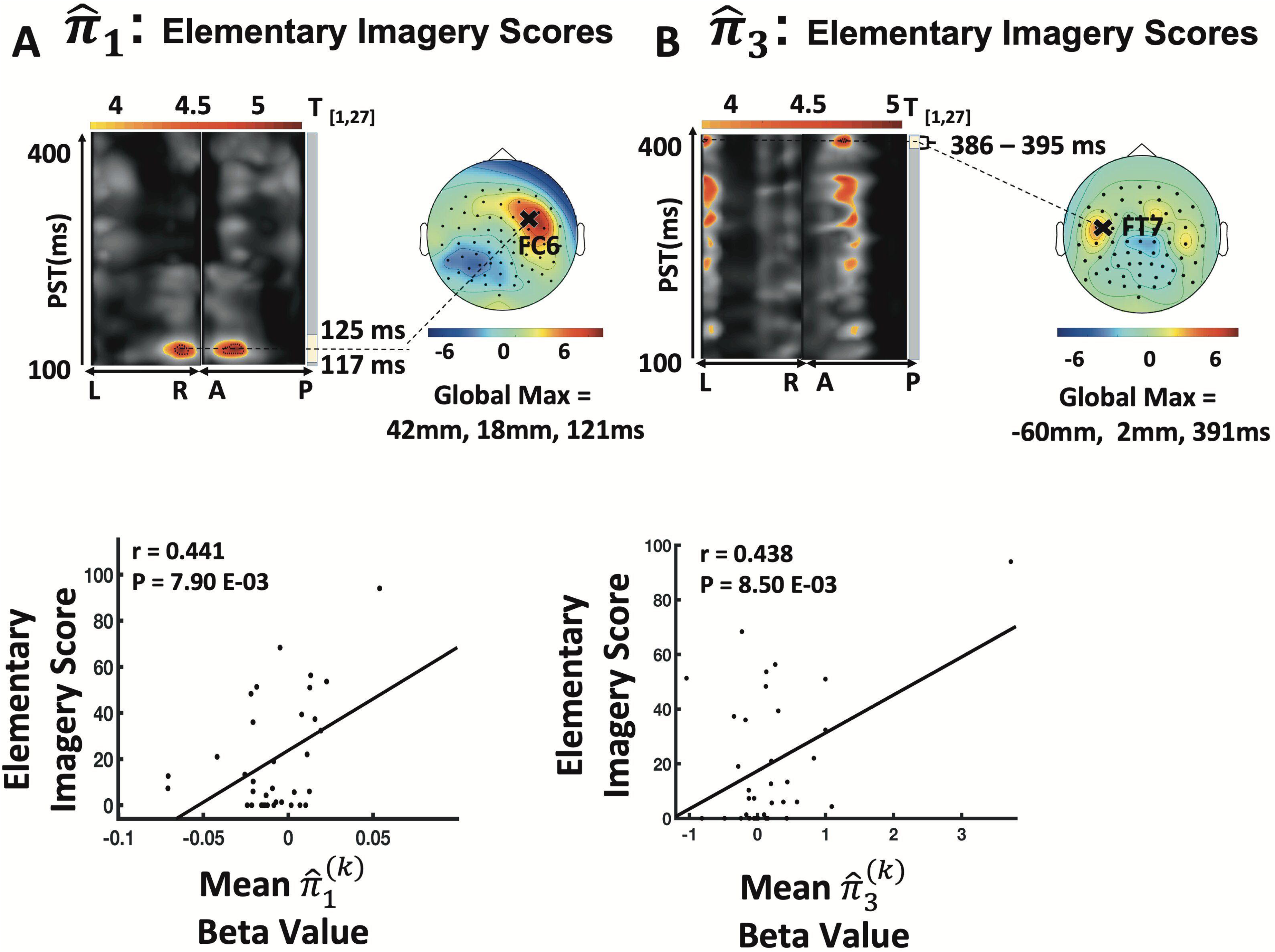
Main effects of precision parameters with 5D-ASC items. (A) Correlations between sensory precision (π^_1_^(*k*)^) and elementary imagery scores (B) Correlations between volatility precision (π^_3_^(*k*)^) and elementary imagery scores. Maximum intensity projections of significant clusters across space ((L) left to (R) right and (A) anterior to (P) posterior) and peri-stimulus time (PST) are shown as topographical maps. Cluster extent is highlighted by the vertical yellow bar. Jet colour mapping displays clusterlevel effects (p*<*0.05, whole-volume family wise error (*FWE*) corrected at the cluster level with a cluster-defining threshold of p*<*0.001) while significant peak effects (p*<*0.05, whole-volume FWE-corrected at the peak level) are denoted with black dotted contours. Scalp maps to the right show cluster effects in jet colour map across a 2-D sensor layout. Scatter plots below illustrate the correlation between scores on ASC scale items and beta values at the global maximum of the T statistic. The correlation coefficients (r) and p-values are reported in the upper left-hand corner. Global maximum coordinates are reported below the scalp maps.

#### Impaired Control and Cognition and Disembodiment

None of the precision signals (sensory (π^_2_^(*k*)^), belief (π^_3_^(*k*)^), or volatility (π^_1_^(*k*)^), showed drug-related differences or main drug effects for impaired control/cognition (π^_2_^(*k*)^: peak 109 ms, *F*_(1,27)_ = 14.67, *p_clus_* = 0.631, sensor P5; π^_1_^(*k*)^peak 121 ms, *F*_(1,27)_ = 16.89, *p_clus_* = 0.562, sensor TP8; π^_2_^(*k*)^: no surviving clusters). No drug differences emerged for disembodiment.

#### Precision-Weighted Prediction Errors

After covariate correction, low-level pwPEs (ε^(*k*)^) showed a lack of drug-related differences. In contrast, high-level pwPEs (ε^(*k*)^) differed markedly between drug groups: an omnibus F-test indicated that ketamine weakened, whereas psilocybin strengthened, pwPE-to-EEG coupling (peak = 309 ms, *F*_(1,27)_ = 31.65, *p_clus_* = 0.001, *p_peak_* = 0.012, sensor: Oz). Follow-up t-tests confirmed this bidirectional pattern, namely reduced correlations under ketamine at 156 ms/posterior electrodes (*T*_(1,27)_ = 5.66, *p_clus_* = 0.030, *p_peak_* = 0.010, sensor: P10) and enhanced correlations under psilocybin at 309 ms/occipital electrodes (*T*_(1,27)_ = 5.89, *p_clus_* = 0.0001, *p_peak_* = 0.006, sensor: Oz).

### Associations with Subjective Experiences

#### Elementary Imagery

ε_3_^(*k*)^-related activity differed by drug, peaking at 316 ms over occipital channels (*F*_(1,27)_ = 30.90, *p_clus_* = 0.002, *p_peak_* = 0.025, sensor: O1). Although both groups showed a positive ε^(*k*)^–imagery correlation, the effect was more pronounced under psilocybin alone at 316 ms (peak = 316 ms, *T*_(1,27)_ = 5.16, *p_clus_* = 0.014, *p_peak_* = 0.030, sensor: O1) (Figure 6B), whereas the ketamine effect did not survive outlier correction.

**Figure 6:**
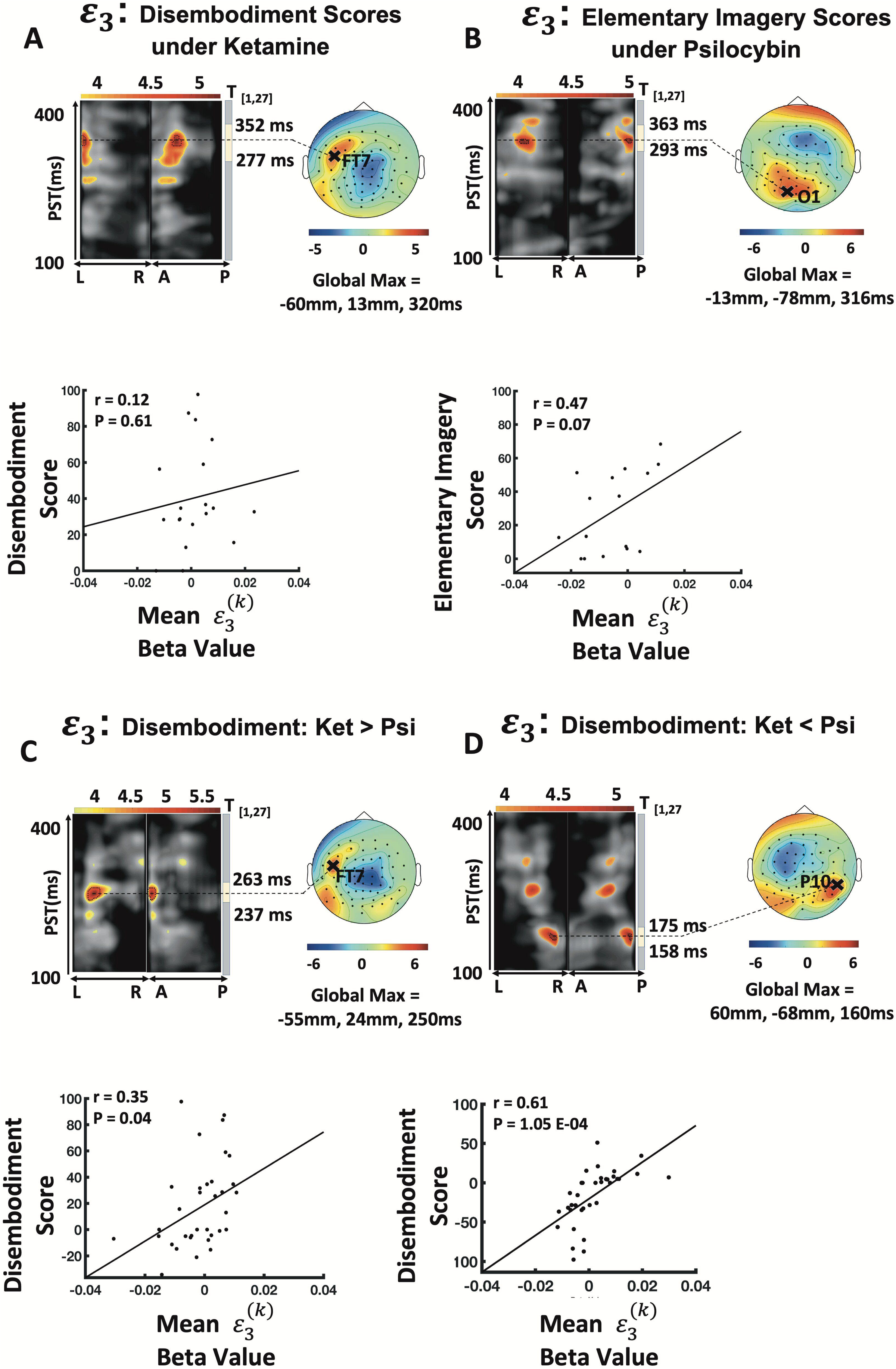
Main effects of precision-weighted prediction error parameters with 5D-ASC items. (A) Correlations between high-level pwPE (ε_3_^(*k*)^) and disembodiment under ketamine (B) Correlations between high-level pwPE (ε_3_^(*k*)^) and elementary imagery under psilocybin (C) (D) Correlations between high-level pwPE (ε_3_^(*k*)^) and ketamine and psilocybin disembodiment scores for (ε_3_^(*k*)^). Maximum intensity projections of significant clusters across space ((L) left to (R) right and (A) anterior to (P) posterior) and peristimulus time (PST) shown as topographical maps. Cluster extent is highlighted by the vertical yellow bar. Jet colour mapping displays cluster-level effects (p*<*0.05, whole-volume family wise error (*FWE*) corrected at the cluster level with a cluster-defining threshold of p*<*0.001) while significant peak effects (p*<*0.05, whole-volume FWE-corrected at the peak level) are denoted with black dotted contours. Scalp maps to the right show cluster effects in jet colour map across a 2-D sensor layout. Scatter plots below illustrate the correlation between scores on Altered States of Consciousness (ASC) [**47**] scale items and beta values at the global maximum of the T statistic. The correlation coefficients (r) and p-values are reported in the upper left-hand corner. Global maximum coordinates are reported below the scalp maps.

#### Disembodiment

High-level pwPEs also tracked disembodiment ratings, with clear drug differences (peak= 250 ms, *F*_(1,27)_ = 35.87, *p_peak_* = 0.010, sensor: FT7). Ketamine showed the stronger ε^(*k*)^–disembodiment coupling at 320 ms (peak = 320 ms, *F*_(1,27)_ = 29.05, *p_peak_* = 0.036, sensor: FT7), but psilocybin exhibited a separate positive correlation at 160 ms (peak = 160 ms, *T*_(1,27)_ = 5.06, *p_peak_* = 0.036, sensor: P10). Two-sample t-tests reflected this bidirectional pattern: ketamine > psilocybin at FT7 (peak = 320 ms, *T*_(1,27)_ = 5.39, *p_clus_* = 0.025, *p_peak_* = 0.018, sensor; FT7; Figure 6A) and psilocybin > ketamine at P10 (peak = 160 ms, *T*_(1,27)_ = 5.25, *p_clus_* = 0.071, *p_peak_* = 0.024, sensor: P10).

#### Impaired Control and Cognition

We did not detect any significant drug differences between ε^(*k*)^ effects and impaired control and cognition in the ketamine and psilocybin groups (ketamine: peak = 293 ms, *T*_(1,27)_ = 4.09, psilocybin: *p_clus_* = 0.140, *p_peak_* = 0.245, sensor: POz; psilocybin: no surviving clusters).

In conclusion, high-level pwPEs are differentially modulated by ketamine and psilocybin: psilocybin enhances, whereas ketamine suppresses, the effects of ε^(*k*)^ and these alterations map onto subjective imagery and disembodiment, but not onto impaired control or cognition.

## 4 Discussion

In this study, we investigated how ketamine and psilocybin modulate the expression of sensory precision, belief precision and hierarchical pwPEs, computational quantities that are required to form an internal model of the world under a predictive coding framework. Our analyses revealed three main findings: Firstly, trial-wise sensory and belief precision signals were significantly weakened under ketamine relative to placebo, whereas psilocybin showed a numerically similar but non-significant reduction; moreover, sensory-precision coupling was weaker under ketamine than psilocybin at 355 ms over posterior sites. Secondly, low-level pwPEs (ε^(*k*)^) were not detectably altered by either drug, but a significant drug × condition interaction revealed that ketamine, not psilocybin, suppressed effects of high-level pwPEs (ε^(*k*)^). Thirdly, correlations with 5D-ASC self-reports indicated differential links between computational signals and subject-experience dimensions, especially elementary imagery and disembodiment.

### 4.1 A ketamine-specific effect: The reduced expression of belief and sensory precision

Factorial analyses showed that both ketamine and psilocybin diminished correlations with Bayes-optimal belief precision (π^^(*k*)^), but only the ketamine effect survived post-hoc comparison with placebo, confirming a ketamine-specific weakening of predicted belief precision. This attenuation could reflect weaker prior under psychotomimetic – mechanism also hypothesized for psychedelics [103] this attenuation likely reflects weaker priors under psychotomimetics. According to the HGF belief update equation (Eq. 1), an imbalance in prior (belief) precision relative to sensory precision shifts the posterior estimates toward the incoming sensory data. Thus, perceptual inference may be driven more strongly by the sensory input.

Post-hoc t-tests further revealed that the drug effect on belief precision was driven principally by ketamine. Complementing this, both ERP and model-based analyses showed that EEG correlates of sensory precision (π^_1_^(*k*)^) were reduced under ketamine peaking at 277 ms in central channels and overlapping with the P3a component in terms of topography. Our dual-level analysis (quantile-binned ERP- and model-based) yielded a consistent picture: Ketamine broadly down-regulated precision signals across the hierarchy: in both single-trial and binned data it significantly attenuated sensory precision (π^_2_^(*k*)^) at ∼280 ms and belief precision (π^_1_^(*k*)^) relative to placebo. By contrast, psilocybin showed minimal impact yet transiently increased early posterior sensory precision (π^_1_^(*k*)^ at 156 ms). The π^_1_^(*k*)^ effect also covaried with the vividness of elementary imagery at 121 ms, with the strongest association under ketamine and a weaker, trend-level effect under psilocybin. This suggests that NMDAR antagonism may amplify low-level sensory processing that, in turn, heightens internally generated imagery [14, 59].

Although higher-level computations are generally assumed to operate at slower temporal scales [58], our early (∼100–200 ms) effects align with previous findings showing rapid neural signatures of belief (π^^(*k*)^) and volatility precision (π^^(*k*)^) between 113–219 ms [25]. In the study by Charlton et al. (2025), high-level precisions were localised in the right STG and left A1 at 254ms. Such rapid engagement of the auditory hierarchy, centred on the STG and bilateral A1, likely reflects specialised adaptation mechanism within auditory prediction circuitry [25].

Collectively, these ketamine-related changes mirror computational mechanisms proposed for the prodrome of psychosis [60, 61, 62, 63]. Aberrantly precise sensory data together with imprecise priors can render lower-level pwPEs salient, fostering delusional explanations [59, 64, 65, 63, 66], but note that also stronger prior beliefs have been found [67, 68] – possibly arising from a compensatory mechanism [69, 70]. Empirical support comes from a reward-learning fMRI study in unmedicated clinical-high-risk individuals, where reduced prior precision and heightened volatility boosted low-level pwPEs and were associated with poorer social and role functioning [71].

Beyond belief-updating, decreased prior precision may contribute to delusions of control in schizophrenia-spectrum disorders (SSDs). Here, imprecise predictions about action outcomes can shift agency attributions to external sources [63, 75]. Ketamine reliably induces such experiences [76] by presumably by disrupting the balance between priors and sensory evidence [77, 78], yet in our data no significant correlations emerged between precision signals and impaired control/cognition, suggesting belief updating mechanisms may be involved (see section 4.2).

Finally, while early theories posited that hallucinations stem from weak priors [79, 80], converging evidence now points to excessively *strong* priors combined with imprecise sensory data [81, 82, 83, 84]. Thus, hallucinations and delusions may arise from opposite imbalances in precision signalling, an idea that future work could test by integrating computational assays with pharmacological interventions modulating NMDAR-plasticity.

### 4.2 Drug-by-condition interactions for higher-level volatility pwPEs

A significant drug × condition interaction emerged for the volatility pwPE or ε_3_^(*k*)^. Relative to placebo, ketamine markedly weakened the correlation between ε_3_^(*k*)^trajectories derived from the ideal Bayesian learner and EEG amplitudes, whereas psilocybin produced no such attenuation. Directly contrasting drug-induced changes (placebo – drug) confirmed this pattern: ε_3_^(*k*)^coupling was significantly lower for ketamine than for psilocybin, peaking at 230 ms over fronto-central electrodes. The binned-ERP contrasts are fully congruent with the single-trial modelling results, with ketamine showing a fronto-central attenuation of ε_3_^(*k*)^-driven ERPs at 223ms. Hence, ketamine selectively dampens higher-level error signalling, in line with previous results [31] and consistent with a tendency to bias learners toward increased learning of environmental volatility. [Psilocybin as serotonergic (5-HT₂A) agonist, can alter unlike ketamine visual perception and higher-level processes [86, 87, 88, 89, 90]. In terms of elementary imagery, although high-level pwPE activity (ε_3_^(*k*)^) predicted the vividness of elementary imagery for both drugs, the effect was markedly stronger under psilocybin at 316 ms over occipital regions. Convergent fMRI studies showed that psilocybin reduced feed-forward connectivity in early visual cortex, thereby amplifying intrinsic V1 activity [91]. Such amplification can evoke canonical geometric or “Klu ver forms” [92, 93, 94]. The associated boost in sensory-precision coupling suggests that the visual system may treat these internally generated visual patterns as if they were high-confidence external input, potentially leading to larger updates of higher-order beliefs. Behaviourally, this may manifest as more vivid autobiographical imagery, even while spatial and temporal working memory deteriorate [95, 96], a profile consistent with REBUS-style relaxation of higher-level priors [39].

A contrasting bidirectional pattern emerged for disembodiment. Ketamine induced larger ε_3_^(*k*)^-disembodiment correlations (peak ≈ 250 ms, fronto-central channels), fitting its well-known dissociative and psychotomimetic profile [97, 98]: NMDA blockade disrupts multisensory integration and bodily self-representation [99, 98]. The resulting “observer stance” may, paradoxically, facilitate therapeutic distancing from maladaptive thoughts [41, 100]. Conversely, psilocybin elicited the stronger correlation at an earlier 160 ms posterior peak, implying a distinct route to altered bodily awareness [101, 102]. Together, these timing and topographic differences highlight how glutamatergic versus serotonergic psychedelics engage partially separate processes to shape visual imagery and the sense of self.

### 4.3 Potential Clinical Implications

Ketamine and psilocybin are routinely employed as pharmacological models of psychosis because they can transiently evoke core schizophrenia-like symptoms - including hallucinations, delusions, thought disorder, and cognitive deficits - in healthy individuals [76, 17, 103]. Early work investigating dissociative anaesthetics such as PCP and ketamine linked these effects to NMDA-receptor antagonism, showing that both positive and negative symptoms could be reproduced via this mechanism. Computational modelling has since refined our understanding by demonstrating that ketamine disrupts hierarchical belief-updating: it reduces correlations with high-level pwPEs derived from an ideal observer model [31] potentially biasing learning towards an exaggerated sense of environmental volatility as observed in early psychosis [104].

Critically, we have observed the same pattern - reduced high-level pwPEs-in individuals at clinical-high risk for psychosis and in patients with first episode psychosis [33, 24] suggesting that the impact of ketamine mimics a key computational deficit underlying the prodrome. Nevertheless, Hauke et al. (2023) found that the magnitude of *low-level* sensory pwPEs (ε_2_^(*k*)^) - not ε_3_^(*k*)^-that distinguished CHR converters from non-converters, underscoring the complementary prognostic value of different hierarchical levels [24]. Mechanistically, abnormalities in low-level PE processing appear to depend on additional mechanisms such as muscarinic receptor signalling [105], indicating that distinct neurochemical pathways may govern learning at different levels of the belief hierarchy. For example, several HGF studies have linked overestimation of environmental volatility (captured at the third level in the hierarchy) to positive symptoms - particularly paranoia – across clinical and sub-clinical samples [52, 66, 104, 24, 106], it should be noted that computational investigations of ε_3_^(*k*)^ did not explicitly assess symptom correlations, and this could be an important future direction.

Taken together, these hierarchical distinctions converge on a broader picture in which aberrant inference permeates multiple representational levels of the cortical hierarchy. Our auditory-domain findings reinforce this view: they show that disruptions in hierarchical prediction-error processing extend beyond higher-order cognition to encompass early sensory prediction mechanisms as well. These observations support predictive coding models of psychosis and underscore the potential of HGF-derived computational markers as neurocognitive markers of prodromal psychosis [104].

The therapeutic effects of psychedelics and psychotomimetics hold promise in a clinical setting. Both ketamine and psilocybin have demonstrated promising antidepressant properties, as shown in randomized control trial data [107, 7, 108]. However, challenges remain with respect to patient-targeted treatment. Heterogeneity of treatment response is a central problem in MDD treatment [109, 110]. Combining computational models with electrophysiological recordings may produce clinically relevant predictions of disease prognosis, based on PE and precision effects. These parameters, which are mechanistically interpretable, could facilitate individualized treatment planning, tailored to a patient’s specific response to a drug’s class, dosage, and interactions (e.g., glutamatergic vs serotonergic psychedelics, psychedelic-assisted vs conventional therapy), aiding in the identification of candidates for specialized treatment regimens in MDD. Our findings indicate these drugs exert complex influences on perceptual processing, which may bear significance for managing psychiatric conditions marked by perturbed sensory and cognitive functions.

### 4.4 Limitations and Future Directions

While our findings provide evidence in support of modulatory effects of s-ketamine and psilocybin on the precisions of predictions and pwPEs, we acknowledge some limitations of this analysis. Firstly, due to the passive nature of the MMN paradigm, we could not fit individual behavioral responses to determine subject-specific trajectories. The interpretation of our results rests on parametric analyses using Bayes-Optimal learning trajectories, which assumes participants are engaging in the optimal minimizing of surprise and learn adaptively across the task sequence. Without subject-specific parameter estimation, interpreting results is more challenging because differences between drug groups may be attributed to varying model suitability. Future research should explore the representation of trajectories using oddball paradigms that necessitate an explicit response from participants.

Secondly, we also recognize that our analysis was also constrained with respect to power, due to small sample sizes (N=19 for the ketamine group and N=16 for psilocybin group), although the within-subject designs increased effective power. Nonetheless, the current results need to be replicated in a study with larger sample sizes.

### 4.5 Conclusions

Taken together, our findings suggest that ketamine and psilocybin have differential effects on sensory learning. Our findings provide valuable insights into the effects of ketamine and psilocybin on optimal Bayesian belief trajectories of higher-level belief precisions and pwPE. These results contribute to understanding the neurobiological mechanisms underlying the cognitive effects of these drugs and highlight the need for further investigation into their therapeutic potentials.

## End Notes

## Acknowledgements

We are grateful for support from the Swiss National Science Foundation (Doc.Mobility P1BSP3-200054 to DJH, Ambizione, PZ00P3-167952 to AOD), the Krembil Foundation (to AOD) and the Swiss Neuromatrix Foundation (to FXV).

## Author Contributions

All authors contributed substantially to this work as outlined below: Concept and design: AS and FXV.

Acquisition, analysis, or interpretation of data: All authors. Drafting of the manuscript: GA and AOD. Critical revision of the manuscript for important intellectual content: All authors. Obtained funding for the original studies: AS, FXV. Supervision: FXV, AOD.

