## Supplemental material for "Ketamine and Psilocybin Differentially Impact Sensory Learning During the Mismatch Negativity"

### Supplementary Information for Ketamine and Psilocybin Differentially Impact Sensory Learning During the Mismatch Negativity

**Shona G. Allohverdi, MSc[1,2], Milad Soltanzadeh, MSc[1,2], André Schmidt, PhD [3], Colleen E. Charlton, MSc [2], Daniel J. Hauke, PhD[4], Povilas Karvelis, PhD[2], Franz X. Vollenweider, PhD †[5], and Andreea O. Diaconescu, PhD †[1,2]**

[1]Institute of Medical Science, Toronto, Canada

[2]Centre for Addiction and Mental Health, Toronto, Canada

[3]Department of Clinical Research (DKF), University of Psychiatric Clinics (UPK), Translational Neurosciences, University of Basel, Switzerland

[4]Hawkes Institute, Department of Computer Science, University College London, London, UK

[5]University Hospital of Psychiatry, Neuropsychopharmacology and Brain Imaging Zurich, Zurich, Switzerland

† Equal contribution

### 1 Materials and Methods

#### 1.1 Participant Inclusion Criteria

Prior to inclusion in the study, all subjects underwent medical evaluations. Their physical health was assessed comprehensively from medical history, clinical examinations, electrocardiography and blood analysis. An absence of a history of drug dependence or present drug abuse was confirmed through urinalysis with a questionnaire on drug consumption. Additionally, subjects were required to confirm that they had refrained from eating breakfast or ingesting caffeine before administration of the drug.

Subjects were selected based on the following inclusion criteria for mental status, each assessed using three different diagnostic checklists:

- DIA-X diagnostic expert system [1]: A computerized, semi-structured interview that algorithmically evaluates DSM-IV and ICD-10 criteria across mood, anxiety, somatoform, and substance-use disorders. In our study, administration took approximately 60–90 minutes, and "no lifetime or current diagnosis" flags were required for inclusion.

- Hopkins Symptom Checklist (SCL-90-R) [2]: 90-item self-report measure assessing the severity of nine symptom dimensions (e.g., depression, anxiety, hostility). Participants scoring below the normative clinical threshold (Global Severity Index < 0.82) were deemed free of significant psychopathology.

- Mini-International Neuropsychiatric Interview (MINI) [3]: A brief (≈ 15 min) structured interview used to confirm the absence of current major psychiatric syndromes, including mood, psychotic, and substance-use disorders. Subjects had to answer "no" to all screening questions.

#### 1.2 Auditory Paradigm

EEG data was recorded during an auditory roving oddball paradigm. Auditory stimuli consisted of trains of seven pure sinusoidal tones ranging in frequency from 500 Hz to 800 Hz in increments of 50 Hz. Trains of given frequency were interrupted by a train of a different frequency. Consequently, a single roving changing tone was established throughout the paradigm. Stimulus trains were presented through Telephonics® TDH-39-P audiometric headphones and generated in E-prime software [50] (TDH-39-P; Maico, Minneapolis, MN). Each tone sequence was unique to each subject and session. Tone duration was set at 70 ms with an inter-stimulus interval of 500 ms. To divert their attention away from the tones, participants performed a visual distracting task in parallel. Whenever a fixation cross changed its luminance, which occurred pseudo-randomly every 2–5 s (not coinciding with auditory changes), subjects had to press a button. One experimental session lasted approximately 15 minutes. Participants were instructed to ignore the tones.

#### 1.3 Results of Visual Distraction Task

Results from the distracting visual vigilance test demonstrated both drugs produced significantly prolonged reaction times ($F(1, 37) = 31.4$, $P<0.0000$, $n^2 = 0.46$) as well as reduced sensitivity indices (*d'*) ($F(1, 37) = 20.8$, $p < 0.00001$, $n^2 = 0.36$)) (see [4]; Table 1).

#### 1.4 Data Processing

Any channels in which over 20% of trials had to be rejected were marked as bad and interpolated for sensor level statistics. S-Ketamine/Placebo group: Bad channels occurred in 5 subjects in placebo session (with 2 (F1,

C2), 5 (Fp1, AF7, Fpz, Fp2, AF8), 2(F1, C2),1 (F2) and 1(F7) channel(s) respectively) with an average of 1211 (SD,201) artefact-free trials 2. Bad channels occurred in 4 subjects in the ketamine session (2 (F1, C2), 1 (FC5), 2 (F1, C2) and 4 (F1) channel(s) respectively). The number of artefact-free trials was significantly less in the placebo session ($p < 0.001$). Psilocybin/Placebo group: Bad channels occurred in 3 subjects in placebo session (with 1 (Fpz), 2 (AF7) and 4 (Fp1, AF7, Fpz, Fp2) channel(s) respectively). Bad channels occurred in 5 subjects in the psilocybin session (2 (FT7, CP1),1 (O1),3 (Fp1, Fpz, Fp2), 1 (O1) and 2 (01, PO8) channel(s) respectively). No significant difference in the number of artifact-free trials between the psilocybin and placebo sessions was found. Similar to Weber and colleagues [5], we employed an ocular movement artifact rejection procedure. All trials that overlapped with the eye blink events were rejected. An additional artefact rejection procedure was applied to remove any problematic trials or channels. Trials with signal across any channel that peaked above ±80 $\mu$V relative to peristimulus baseline were removed.

#### 1.5 General Linear Model

##### 1.5.1 First level GLMs

Vectors of the Bayes optimal model-based pwPE ($\varepsilon_2^{(k)}, \varepsilon_3^{(k)}$) and precision ($\hat{\pi}_1^{(k)}, \hat{\pi}_2^{(k)}$ and $\hat{\pi}_3^{(k)}$) parameters were used as regressors in a multiple regression model of trial-wise EEG signals for each subject in each session. Although the same regressors were used for all subjects, entries corresponding to rejected trials during EEG preprocessing were excluded to maintain consistency in regressor length across subjects. Regressors were not orthogonalized. Statistical maps of the resulting beta coefficients were corrected for multiple comparisons over the full time-sensor volume via Gaussian random field theory [6]. These first-level beta images then served as inputs to group-level inference, with all reported effects surviving either peak- or cluster-level family-wise error (FWE) correction.

1. Precision-weighted prediction errors:

$$Y = \beta_0 + X\beta + \epsilon$$

$$Y = [y_1 \ldots y_n]' \qquad X = [\epsilon_2, \epsilon_3] \qquad \beta = [\beta_1 \beta_2]'$$

2. Precisions:

$$Y = \beta_0 + X\beta + \epsilon$$

$$Y = [y_1 \ldots y_n]' \qquad X = [\hat{\pi}_1, \hat{\pi}_2, \hat{\pi}_3] \qquad \beta = [\beta_1 \beta_2 \beta_3]'$$

$n$ = number of trials, $\epsilon$ = error

#### 1.6 3D Scalp–Time EEG Image Construction and Pseudo-Condition Difference-Wave Computation

Following pre-processing and artefact correction as described above, pre-processed single-trial EEG data were converted to 3-dimensional volumes with two spatial dimensions (anterior-to-posterior and left-to-right directions of the scalp surface) and a temporal dimension (peri-stimulus time). These scalp × time 3D images (voxel size: 4.2mm x 5.4 mm x 3.9 ms) were subjected to statistical analysis using the GLM in an analogous fashion to functional magnetic resonance imaging [7]. This is a well-established method for statistical analysis of scalp EEG data using the SPM software [8]. Using this approach, we pursued single trial analyses where optimal Bayesian computational model trajectories provided trial-wise regressors that were used to explain EEG amplitudes.

The generation of the difference waveform was accomplished through an initial computation of grand averaged Event-Related Potentials (ERPs) based on two distinct pseudo-conditions obtained by sorting and subsequently averaging trials corresponding to the the top and bottom $20^{th}$ percentiles of each computational quantity's magnitude. Following this, the difference waveform was derived by subtracting those two waveforms.

#### 1.7 Psychometric Assessment of Altered States

A modified form of the 5D-ASC was used to explore the cognitive and perceptual effects mediated by both drugs. From five original dimensions, we investigate three that measure the subjective effects: (i) Oceanic Boundlessness (OB) which is comprised of the subscales of blissful state, spiritual experience, insightfulness, disembodiment and experience of unity, (ii) Visionary Restructuralization (VR) with subscales elementary imagery, complex imagery, audio-visual synesthesia, changed meaning of percepts and auditory alterations, and (iii) Dread of Ego Dissolution (DED) with subscales of anxiety and impaired control and cognition.

In our study, we focused on three out of the eleven subscales — disembodiment, elementary imagery, and impaired control and cognition — due to their relevance in this study. These subscales were distinctively affected by the two drugs under investigation, as outlined by [4]. Consequently, they were chosen as covariates in the GLM to account for their differential impact.

Based on the samples included in our analysis, we re-examined drug-related differences on ASC-R scores. As reported by Schmidt and colleagues [4], at the doses tested in this study, ketamine and psilocybin similarly affected the global ASC score, showing changes in derealization and depersonalization, mood shifts, cognitive deficits, and sensory distortions. This was indicated by a significant main effect for treatment ($F_{(1,37)} = 69.5$, $p < 0.000001$, $\eta^2 = 0.65$; Table 4).

In contrast to psilocybin, ketamine resulted in significantly higher scores in depersonalization, as measured by disembodiment ($p = 0.0006$), and in cognitive impairments, as measured by impaired control and cognition ($p = 0.01$) (see 4). For additional analyses of the ASC scores, please refer to the original paper by Schmidt and colleagues [4].

### 2 Supplementary Tables

**Table S1. Mean Descriptive Characteristics of subjects by drug group**

| **Characteristic** | **Ketamine(N=19)** | **Psilocybin(N=16)** | ***P*-value** |
|---|---|---|---|
| Sex (M/F) | 13/6 | 11/5 | 0.984 |
| Age (Mean years) [SD] | 26 [5.36] | 23 [2.41] | 0.085 |
| Weight (kg) [SD] | 72.94 [12.44] | 68.88 [12.28] | 0.317 |

Supplementary Table 1: Mean descriptive characteristics of subjects in study groups

**Table S2. Mean Trial Statistics [SD]**

| | Ketamine | | | Psilocybin | | |
|---|---|---|---|---|---|---|
| | **Placebo** | **Drug** | Group Difference | **Placebo** | **Drug** | Group Difference |
| Number of eyeblink trials rejected | 236[106] | 180[120] | $p < 0.05$ | 218[58] | 197[95] | $p = 0.380$ |
| Number of artefact-free trials | 1211[201] | 1466[210] | $p < 0.001$ | 1305[169] | 1232[241] | $p = 0.332$ |

Supplementary Table 2: Mean and, in brackets, the standard deviation of the number of eyeblink trials rejected and number of remaining trials

**Table S3. Mean Subject-Specific Perceptual Parameter Settings**

| Parameter | Prior mean | Prior variance | Posterior mean | SD |
|---|---|---|---|---|
| κ | 1 | 0 | - | - |
| ω | −6 | 25 | -10.06 | 0.28 |
| ϑ | 0.05 | 0.088 | 0.05 | 0 |
| $\mu_3^{(k=0)}$ | 1 | 0 | 1 | 0 |
| $\sigma_3^{(k=0)}$ | log(0.1) (≈ −2.303) | 1 | 0.10 | 0 |
| $\mu_{\{2ij\}}^{(k=0)}$ | logit(1/49) (≈ −3.871) | 0 | - | - |
| $\sigma_{\{2ij\}}^{(k=0)}$ | log(1) (= 0) | 0 | - | - |

Supplementary Table 3: Priors on HGF perceptual parameters and starting values. All parameters and starting values were fixed (i.e., not estimated) except for the tonic learning rates $\omega$ and $\vartheta$, and the starting value of the high-level uncertainty $\sigma_3^{(0)}$. For these three parameters, we also report posterior group means and standard deviations. There were no significant differences between the parameters in the placebo and the drug conditions.

| **Table S4. Mean 5D-ASC-R Scores [SD]** | | | | | | |
|---|---|---|---|---|---|---|
| Subscale | Ketamine group (Placebo) (N=19) | Psilocybin group (Placebo) (N=16) | *p*-value | Ketamine group (Drug) (N=19) | Psilocybin group (Drug) (N=16) | *p*-value |
| **Global ASC** | | | | | | |
| | 0.68 [1.60] | 0.41[0.54] | 0.51 | 22.33[16.50] | 19.06[13.41] | 0.52 |
| **Oceanic Boundlessness (OB)** | | | | | | |
| Disembodiment | 0.35[0.82] | 0.37[0.63] | 0.92 | 40.35[28.37] | 11.80[13.70] | **0.0006**** |
| Experience of Unity | 0.62[1.69] | 0.20[0.50] | 0.42 | 29.4[26.18] | 17.41[18.94] | 0.13 |
| Spiritual Experience | 0.60[1.59] | 0.35[0.51] | 0.54 | 13.10[18.44] | 8.50[13.86] | 0.41 |
| Blissful state | 0.60[1.40] | 0.54[0.93] | 0.97 | 25.67[18.65] | 26.40[20.64] | 0.91 |
| Insightfulness | 0.44[1.14] | 0.40[0.68] | 0.89 | 17.82[25.09] | 17.90[26.44] | 0.99 |
| **Visionary Restructuralization (VR)** | | | | | | |
| Complex Imagery | 0.60[1.53] | 0.35[0.65] | 0.54 | 17.81[28.47] | 21.00[21.32] | 0.71 |
| Elementary Imagery | 0.84[1.89] | 0.25[0.52] | 0.20 | 13.60[22.89] | 28.00[24.40] | 0.08 |
| AV synesthesia | 1.60[4.73] | 0.67[1.34] | 0.43 | 29.78[29.45] | 28.21[32.16] | 0.88 |
| Changed Meaning of Percepts | 0.74[1.84] | 0.31[0.52] | 0.35 | 20.12[23.10] | 30.35[21.32] | 0.18 |
| **Dread of Ego Dissolution (DED)** | | | | | | |
| Impaired control and cognition | 0.80[1.55] | 0.76[1.36] | 0.94 | 30.60[16.42] | 16.31[15.38] | **0.01*** |
| Anxiety | 0.42[1.02] | 0.32[0.53] | 0.72 | 7.51[10.35] | 3.90[9.45] | 0.28 |

Supplementary Table 4: Mean 5D-ASC-R scores by drug group and condition. *Note*: **p < 0.001 * p < 0.05

| Table S5. Main binned ERP results for all 5 computational quantities | | | | | | |
|---|---|---|---|---|---|---|
| $\hat{\boldsymbol{\pi}}1$ | Contrast | Cluster extent k (voxel) | Cluster PFWE | Peak T | Peak PFWE | Peak coordinates (x mm, y mm, t ms) |
| Ketamine (N=19) | Placebo > Ketamine | 1 372 | 0.0001 | 5.00 | 0.11 | -4, 24, 281 |
| | Ketamine > Placebo | 3 | 0.611 | 3.80 | 0.560 | −21, −78, 285 |
| Psilocybin (N=16) | Placebo < Psilocybin | 177 | 0.086 | 6.70 | 0.028 | 38, 2, 156 |
| | Placebo > Psilocybin | 17 | 0.985 | 4.17 | 0.590 | −47, −14, 203 |
| $\hat{\boldsymbol{\pi}}_2$ | Placebo > Ketamine | 57 | 0.320 | 4.47 | 0.239 | −26, −19, 172 |
| Ketamine (N=19) | Ketamine > Placebo | 3 | 0.596 | 3.48 | 0.607 | −51, −62, 258 |
| | Placebo < Psilocybin | — | — | — | — | — |
| Psilocybin (N=16) | Placebo > Psilocybin | — | — | — | — | — |
| $\hat{\boldsymbol{\pi}}_3$ | Placebo > Ketamine | 90 | 0.239 | 4.81 | 0.145 | 55, −46, 176 |
| Ketamine (N=19) | Ketamine > Placebo | 13 | 0.516 | 3.89 | 0.495 | −13, − 3, 176 |
| | Placebo < Psilocybin | 15 | 0.591 | 4.20 | 0.552 | −13, 68, 312 |
| Psilocybin (N=16) | Placebo > Psilocybin | 6 | 0.680 | 4.20 | 0.554 | −51, 40, 395 |
| $\boldsymbol{\varepsilon}2$ | Placebo > Ketamine | 268 | 0.070 | 4.98 | 0.105 | 8, 29, 223 |
| Ketamine (N=19) | Ketamine > Placebo | 1 | 0.945 | 3.78 | 0.904 | 21, 47, 207 |
| | Placebo < Psilocybin | — | — | — | — | — |
| Psilocybin (N=16) | Placebo > Psilocybin | — | — | — | — | — |

| | | | | | | |
|---|---|---|---|---|---|---|
| $\boldsymbol{\varepsilon_3}$ | Placebo > Ketamine | 108 | 0.203 | 5.07 | 0.197 | −55, −12, 172 |
| Ketamine (N=19) | Ketamine > Placebo | 333 | 0.041 | 5.05 | 0.100 | −4, 18, 168 |
| | Placebo < Psilocybin | 4 | 0.622 | 4.20 | 0.650 | −51, −14, 234 |
| Psilocybin (N=16) | Placebo > Psilocybin | — | — | — | — | — |

Supplementary Table 5: Main binned ERP results based on precision and precision-weighted PE binned data. Significant clusters (p<0.05) highlighted in bold. “—” indicates that no suprathreshold clusters were detected for the respective contrast.

| **Table S6. Main effects of epsilon design ($\varepsilon_2^{(k)}, \varepsilon_3^{(k)}$)** | | | | | | | |
|---|---|---|---|---|---|---|---|
| Computational Parameter | Cluster | $k_E$ | Cluster significance ($P_{FWE}$) | Peak significance ($P_{FWE}$) | $F_{66}$ | $Z\equiv$ | Peak location (mm,mm,ms) |
| $\varepsilon_2$ | 1 | 7690 | **1.95E-12** | **1.83E-11** | 111.6<br>6 | Inf | 13, 8, 176 |
| | 2 | 3216 | **2.89E-07** | **1.24E-10** | 101.0<br>0 | 7.71 | -60, -62, 168 |
| | 3 | 378 | **0.028** | **0.028** | 22.50 | 4.23 | 13, 13, 395 |
| | 4 | 53 | 0.335 | 0.095 | 19.13 | 3.92 | -60, -62, 398 |
| | 5 | 118 | 0.179 | 0.137 | 18.38 | 3.84 | 17, -30, 336 |
| $\varepsilon_3$ | 1 | 2378 | **1.11E-05** | **7.56E-09** | 79.61 | 7.11 | 60, -62, 164 |
| | 2 | 4030 | **9.36E-08** | **1.31E-07** | 66.69 | 6.66 | 8, 29, 164 |
| | 3 | 2116 | **2.60E-05** | **8.11E-07** | 59.03 | 6.36 | -4, -3, 250 |
| | 4 | 1891 | **5.54E-05** | **4.25E-05** | 43.75 | 5.65 | 0, -3, 398 |
| | 5 | 733 | **0.00530** | **1.48E-04** | 39.31 | 5.41 | 60, -62, 398 |
| | 6 | 476 | **0.0191** | **3.90E-04** | 35.96 | 5.21 | -60, -62, 398 |
| | 7 | 524 | **0.0148** | **5.62E-04** | 34.71 | 5.13 | -60, -46, 242 |
| | 8 | 108 | 0.197 | **0.0054** | 26.34 | 4.55 | 8, 18, 102 |
| | 9 | 14 | 0.475 | 0.315 | 14.48 | 3.42 | 0, -89, 262 |
| | 10 | 17 | 0.457 | 0.382 | 13.78 | 3.34 | 60, -19, 113 |
| | 11 | 7 | 0.522 | 0.555 | 12.27 | 3.14 | 60, 29, 379 |

Supplementary Table 6: Main effects of precision-weighted PEs ( $\varepsilon_2^{(k)}$ , $\varepsilon_3^{(k)}$ ). Significant clusters ($p$<0.05) highlighted in bold.

| **Table S7. Main effects of precision design** $(\hat{\pi}_1^{(k)}, \hat{\pi}_2^{(k)}, \hat{\pi}_3^{(k)})$ | | | | | | | |
|---|---|---|---|---|---|---|---|
| Computational Parameter | Cluster | $k_E$ | Cluster significance ($P_{FWE}$) | Peak significance ($P_{FWE}$) | $F_{66}$ | $Z \equiv$ | Peak location (mm,mm,ms) |
| $\hat{\pi}_1$ | 1 | 6192 | **5.87E-10** | **0.000** | 128.54 | Inf | 8, 24, 176 |
| | 2 | 3552 | **4.12E-07** | **8.33E-12** | 115.37 | Inf | 60, -57, 148 |
| | 3 | 4013 | **1.19E-07** | **4.88E-07** | 61.05 | 6.45 | -8, -9, 254 |
| | 4 | 2580 | **6.81E-06** | **5.13E-05** | 43.01 | 5.62 | 0, 2, 398 |
| | 5 | 826 | **0.0036** | **8.97E-05** | 32.02 | 4.96 | -60, -57, 379 |
| | 6 | 186 | 0.112 | **0.012** | 20.29 | 4.03 | -60, -41, 242 |
| | 7 | 2 | 0.563 | 0.437 | 12.09 | 3.12 | 0, -89, 289 |
| $\hat{\pi}_2$ | 1 | 1224 | **0.001** | **2.69E-06** | 54.44 | 6.17 | 13, 2, 184 |
| | 2 | 494 | **0.016** | **0.030** | 22.13 | 4.20 | 42, 13, 141 |
| | 3 | 91 | 0.228 | **0.084** | 18.96 | 3.90 | -60, -57, 176 |
| | 4 | 61 | 0.298 | **0.091** | 18.70 | 3.88 | -26, -9, 125 |
| | 5 | 108 | 0.197 | 0.129 | 17.61 | 3.77 | -60, -41, 133 |
| | 6 | 61 | 0.298 | 0.154 | 17.04 | 3.71 | 21, 8, 398 |
| | 7 | 20 | 0.455 | 0.432 | 13.48 | 3.30 | 8, -84, 168 |
| | 8 | 1 | 0.603 | 0.606 | 12.03 | 3.11 | 55, -25, 141 |
| $\hat{\pi}_3$ | 1 | 4454 | **2.10E-09** | **8.90E-07** | 59.55 | 6.39 | 8, 13, 102 |
| | 2 | 1720 | **2.67E-05** | **1.05E-06** | 58.87 | 6.36 | 60, -62, 113 |
| | 3 | 14 | 0.525 | **0.032** | 22.40 | 4.22 | 0, -89, 102 |
| | 4 | 47 | 0.353 | 0.213 | 16.43 | 3.64 | 64, 8, 324 |
| | 5 | 11 | 0.548 | 0.382 | 14.39 | 3.41 | 64, -3, 168 |
| | 6 | 25 | 0.455 | 0.384 | 14.37 | 3.41 | 64, 8, 258 |
| | 7 | 5 | 0.603 | 0.459 | 13.68 | 3.32 | -26, -25, 324 |
| | 8 | 1 | 0.655 | 0.671 | 11.96 | 3.10 | 21, -78, 102 |

Supplementary Table 7: Main effects of precisions $(\hat{\pi}_1^{(k)}, \hat{\pi}_2^{(k)}, \hat{\pi}_3^{(k)})$. Significant clusters ($p$<0.05) highlighted in bold.

### 3 Outlier Screening and ANCOVA results: ( $\hat{\pi}_3^{(k)}$). Elementary Imagery Scores

To identify statistical outliers within the dataset, the interquartile range (IQR) method was employed. This approach defines outliers as data points that fall below the lower bound (Q1 – 1.5 × IQR) or above the upper bound (Q3 + 1.5 × IQR), where Q1 and Q3 represent the 25th and 75th percentiles, respectively. The IQR, calculated as the difference between Q3 and Q1, captures the middle 50% of the data and serves as a robust measure of variability. This method was selected due to its simplicity and effectiveness in detecting extreme values without making assumptions about the underlying data distribution.

#### 3.1 Outlier Detection and Removal

Using the 1.5×IQR rule, we flagged two extreme values, one β-coefficient of 3.724 (upper IQR bound = 1.08) and one imagery score of 94 (upper IQR bound = 91.67), and removed them from the dataset.

In the original ANCOVA (N=28), ketamine significantly altered the relationship between high-level volatility pwPEs ($\varepsilon_3^{(k)}$) and imagery ratings, revealing a fronto-central cluster peaking at –60 mm/2 mm and 391 ms ($T_{27}$ = 5.10, $p_{\mathrm{FWE}}$ = 0.037). However, once the outliers were excluded (N=27), this effect vanished: the same contrast yielded only a small cluster at 21 mm/50 mm/207 ms that did not survive correction ($T_{26}$ = 3.54, $p_{\mathrm{FWE}}$ = 0.613). These findings indicate that the initial ketamine–imagery coupling was driven by a few extreme data points.

### 4 Supplementary Figures

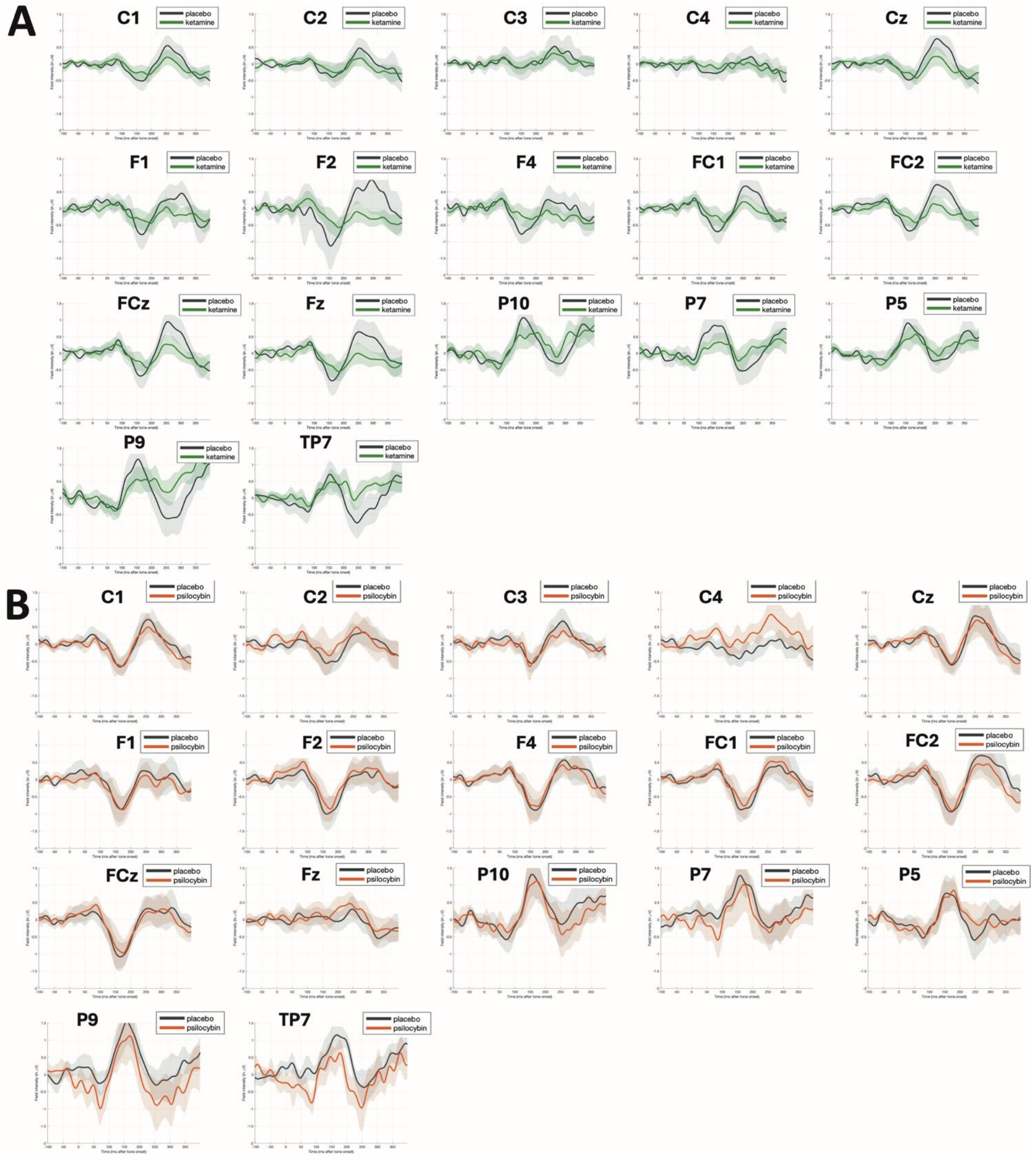

**Supplementary Figure 1**: **Trial-binned ERP difference waves reveal widespread attenuation of sensory-precision signalling under ketamine and psilocybin ( $\hat{\pi}_1^{(k)}$).** Each subplot depicts the high-minus-low quantile difference waveform for the sensory-precision parameter at the indicated scalp electrode (mean ± 1 SEM). Trials were ranked within participant by $\hat{\pi}_1^{(k)}$ estimates from a three-level Hierarchical Gaussian Filter and the upper and lower 20 % were averaged before subtraction. In the upper plot (A) black traces correspond to placebo; green traces to ketamine. In the lower plot (B) black traces correspond to placebo; red traces to psilocybin. Subplots are ordered with the following rows: row 1, central (C1, C2, C3, C4, Cz); row 2, frontal (F1, F2, F4, FC1, FC2); row 3, frontal, temporal–parietal (FCz, Fz, P10, P7, P8); row 4, temporal, posterior/occipital (P9, TP7). Across the scalp, ketamine markedly reduced sensory-precision responses relative to placebo ($T_{(1, 18)}$ = 5.00, $p$<0.001). Psilocybin exhibited the opposite pattern, selectively augmenting sensory-precision-related activity over posterior-central channels between 150 – 300 ms ($T_{(1, 15)}$ = 6.70, $p$ = 0.028).

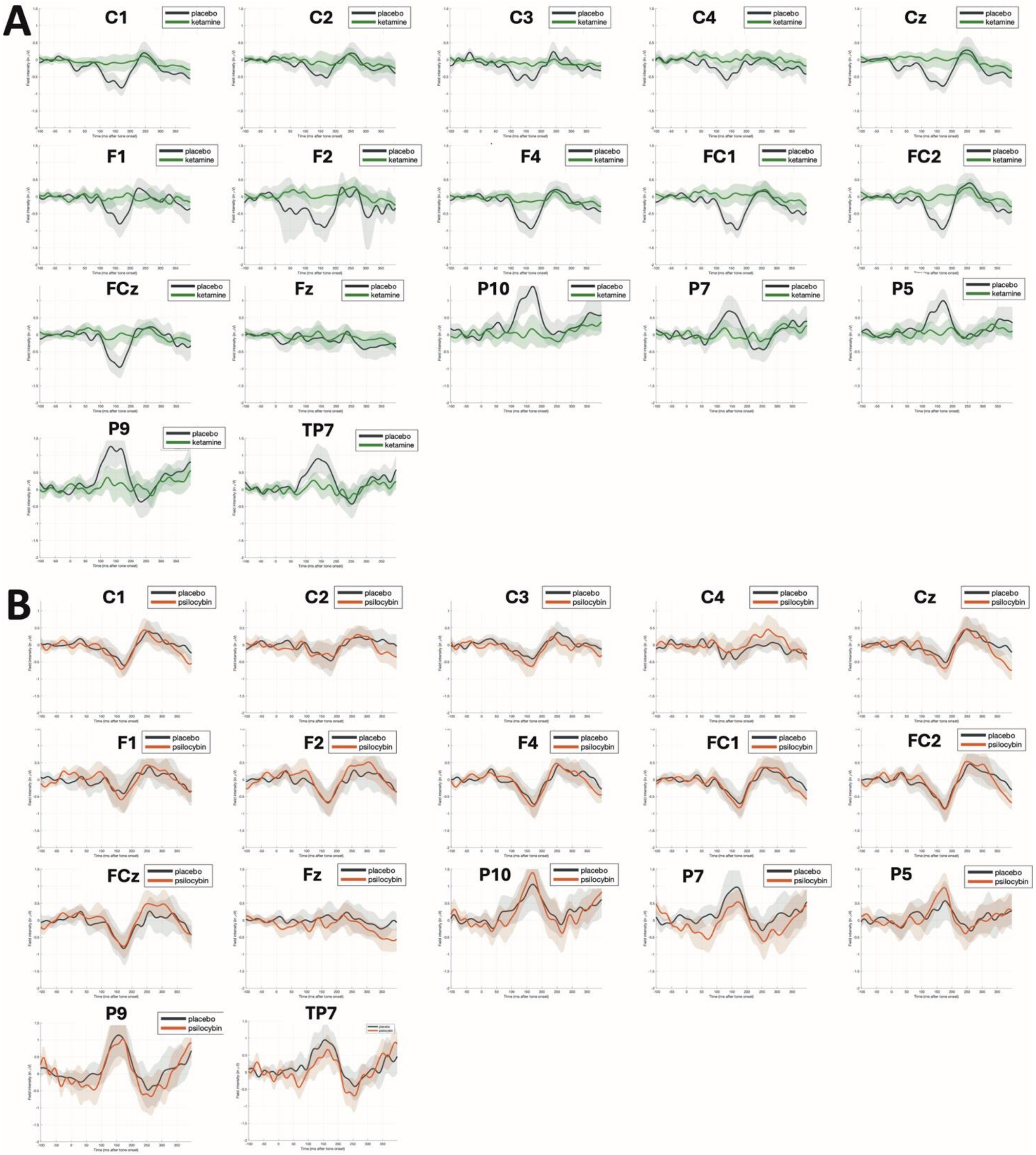
A
C1
C2
C3
C4
Cz
F1
F2
F4
FC1
FC2
FCz
Fz
P10
P7
P5
P9
TP7
placebo
ketamine
B
C1
C2
C3
C4
Cz
F1
F2
F4
FC1
FC2
FCz
Fz
P10
P7
P5
P9
TP7
placebo
psilocybin

**Supplementary Figure 2: Trial-binned ERP difference waves reveal widespread attenuation of volatility precision-weighted prediction error signalling under ketamine and psilocybin ( $\varepsilon_3^{(k)}$).** Each subplot depicts the high-minus-low quantile difference waveform for the volatility precision-weighted PE parameter at the indicated scalp electrode (mean ± 1 SEM). Trials were ranked within participant by $\varepsilon_3^{(k)}$ estimates from a three-level Hierarchical Gaussian Filter and the upper and lower 20 % were averaged before subtraction. In the upper plot (A) black traces correspond to placebo, green traces to ketamine. In the lower plot (B) black traces correspond to placebo, red traces to psilocybin. Subplots are ordered with the following rows: row 1, central (C1, C2, C3, C4, Cz); row 2, frontal (F1, F2, F4, FC1, FC2); row 3, frontal, temporal–parietal (FCz, Fz, P10, P7, P8); row 4, temporal, posterior/occipital (P9, TP7). Across the scalp, ketamine produced a significant fronto-central attenuation of high-level volatility-precision-weighted prediction-error signalling ($T(1, 18) = 5.05$, $p = 0.04$). Psilocybin did not elicit any suprathreshold modulation of $\varepsilon_3^{(k)}$ -binned responses.

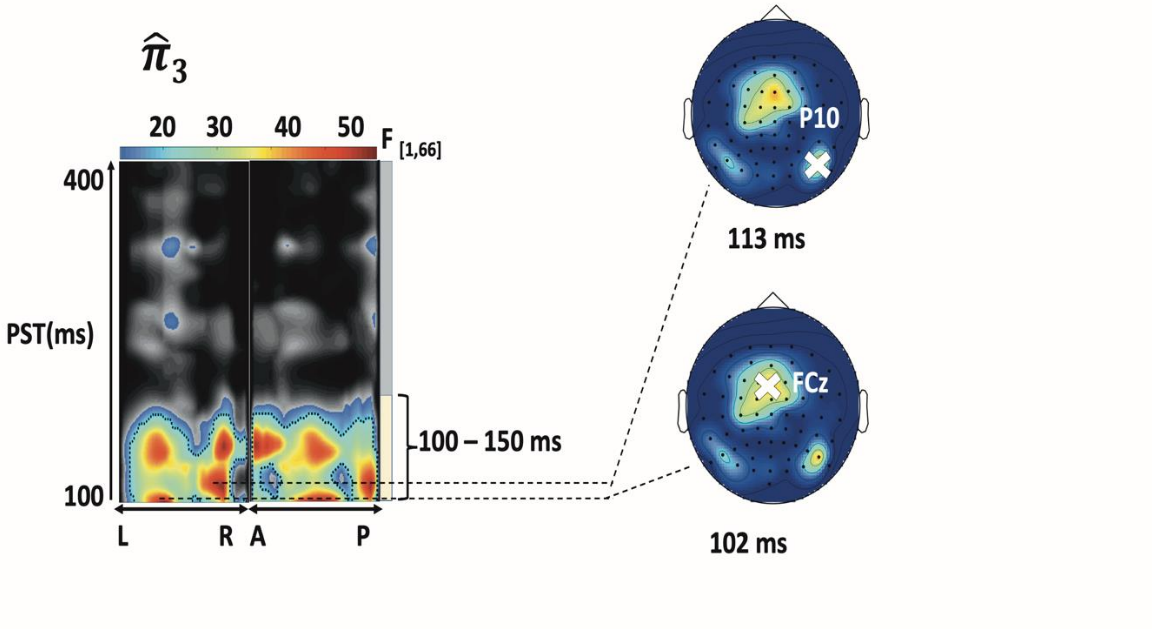


**Supplementary Figure 3: Main effect of ( $\hat{\pi}_3^{(k)}$).** Maximum intensity projections for significant clusters of across cortical space (left to right and anterior to posterior) shown in topographical maps. Cluster extents are highlighted by vertical yellow bar. Jet colour mapping displays cluster-level effects (p<0.05, whole volume family wise error (*FWE*) corrected at the cluster level with a cluster-defining threshold of p<0.001) while significant peak effects (p<0.05, whole-volume FWE-corrected at the peak level with a cluster-defining threshold of p<0.05) are denoted with black dotted contours. **Right** Scalp maps to the right show cluster effects in jet colour map across a 2-D sensor layout.
