## Supplementary figures and images for "Ketamine and Psilocybin Differentially Impact Sensory Learning During the Mismatch Negativity"

### Supplemental figure 1

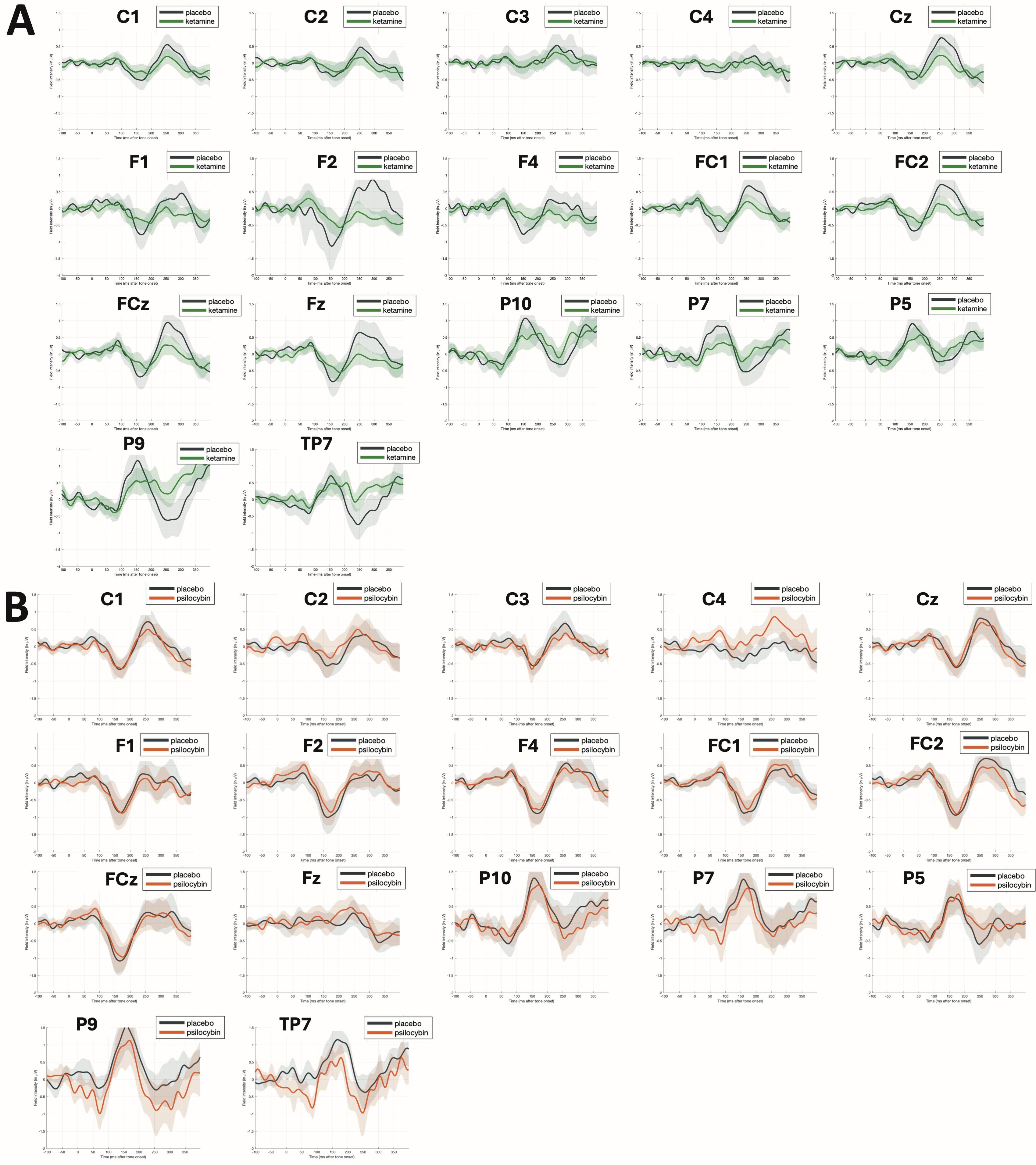

### Supplemental figure 2

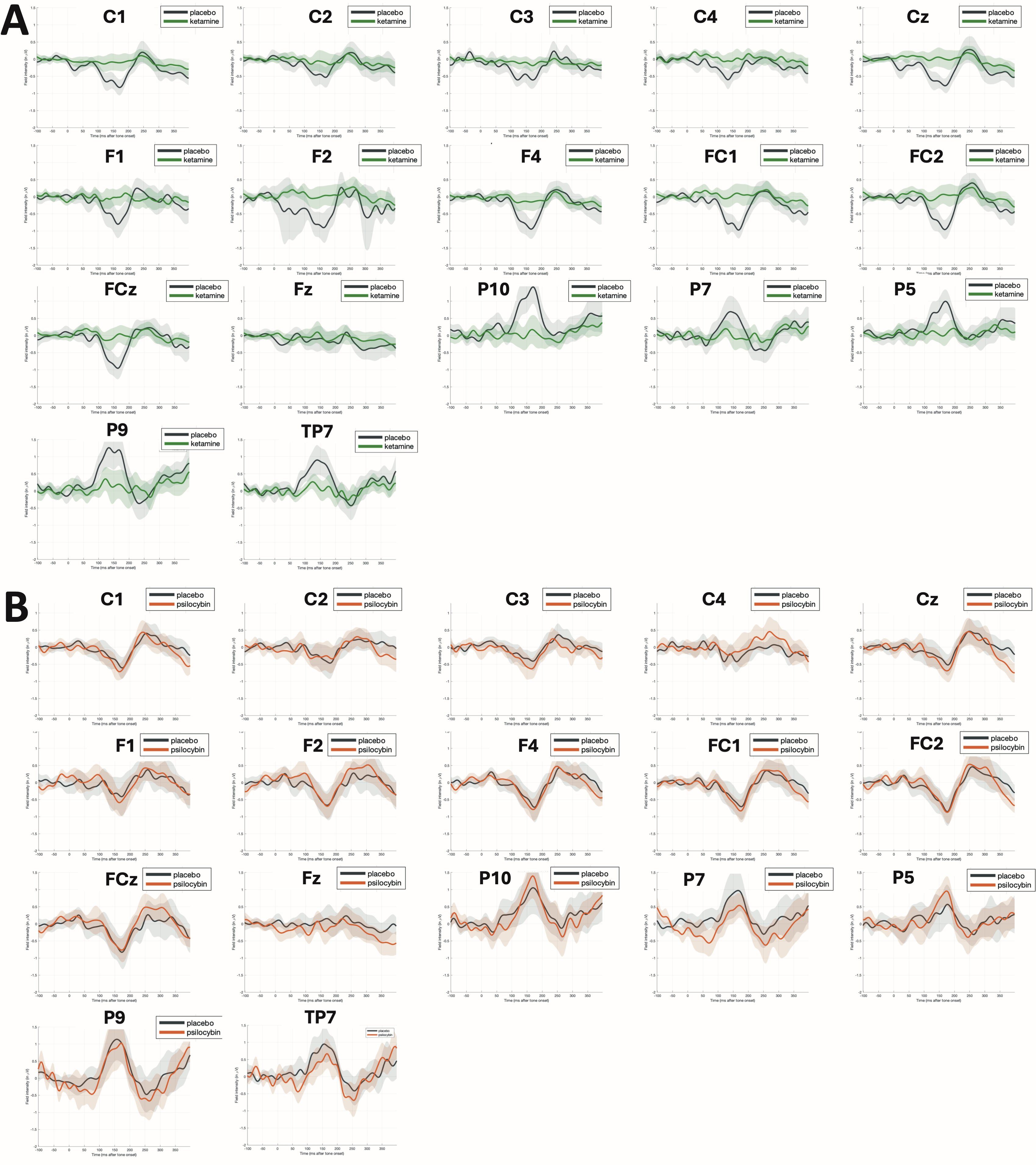

### Supplemental figure 3

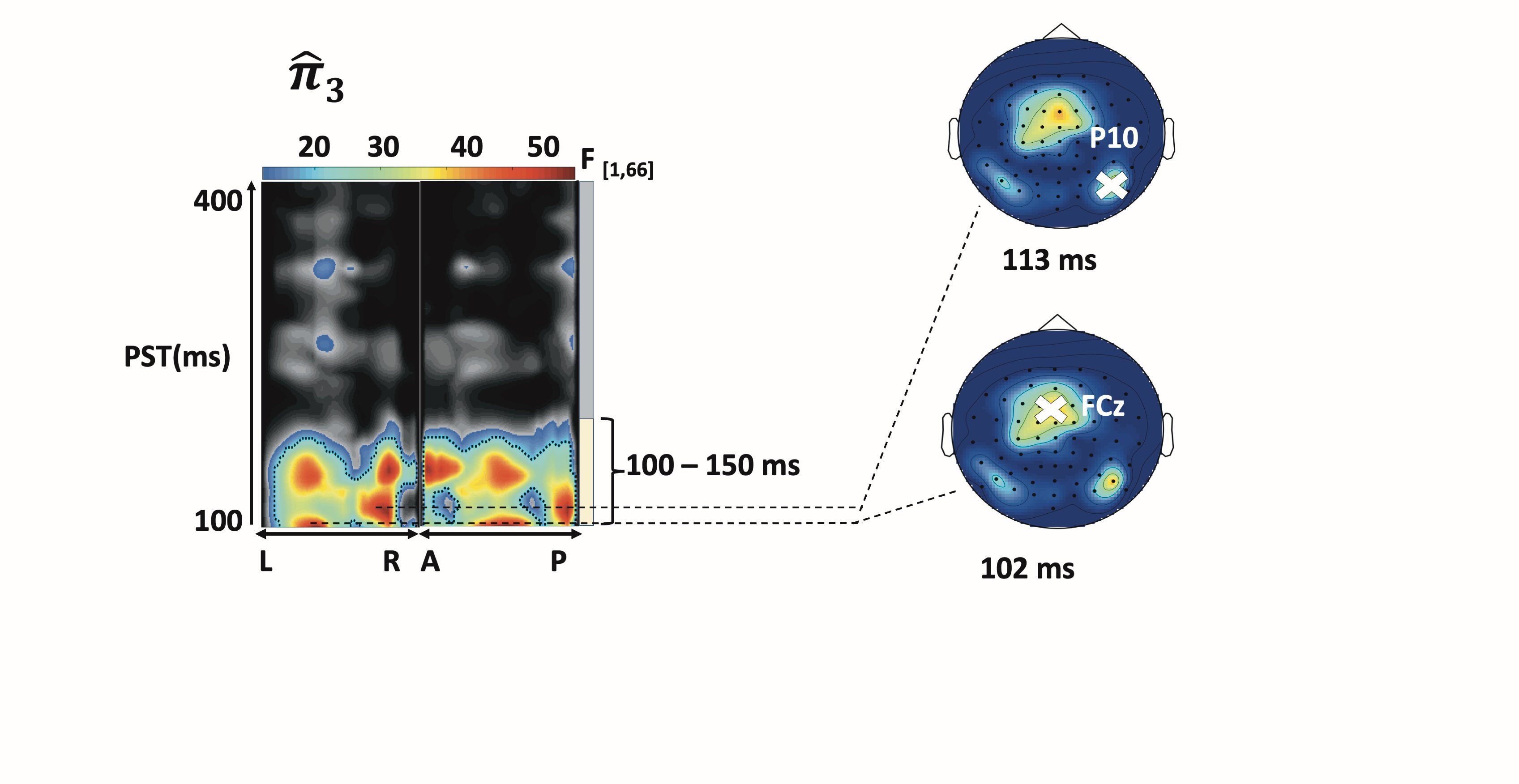
